# Negative selection on baboon admixture is strongest on chromosome X

**DOI:** 10.1101/2024.12.18.629079

**Authors:** Erik F. Sørensen, Garrett Hellenthal, Kasper Munch

**Author notes:** Corresponding Author: Kasper Munch.

## Abstract

The six admixing baboon species offer a natural experiment to study negative selection on admixture and potentially genetic incompatibility in species with divergence resembling that between anatomically modern and extinct human subspecies such as Neanderthals. We analyze 156 high-coverage genomes sampled from seven olive and four yellow baboon populations shaped by admixture in two different reticulation events. In Tanzania, olive and yellow baboons admix despite 1.3 million years of divergence. In Ethiopia, olive baboons have invaded and displaced an ancient Hamadryas-like population separated by 0.6 million years, thus mirroring the displacement of Neanderthals by modern humans. Analyzing local ancestry across whole genome, we find evidence of negative selection on minor parent ancestry in Tanzanian yellow and olive baboon populations reaching far behind the hybrid zone and in Ethiopian olive baboons. We find that evidence of selection against admixture is up to seven times stronger on the X chromosome. The proportion of minor parent ancestry is substantially higher on the X chromosome in Ethiopian olive and yellow baboon populations, which displaced the populations now representing their minor parent ancestry. This additional MPA is concentrated in a few genomic regions with high frequency. This indicates that this original ancestry was retained by negative selection on the invading ancestry, suggesting that these loci play a role in emerging reproductive barriers in the Papio as well as the Homo genus.

## Introduction

Speciation is often a gradual process in which populations diverge due to geographic or ecological isolation. This isolation fosters differentiation as populations accumulate unique mutations and experience distinct selective pressures and genetic drift[1–4]. In cases of prolonged isolation, genetic incompatibilities can emerge[5–7], eventually leading to complete reproductive isolation, thus preventing further admixture[8–10]. Many mutations are nearly neutral[11–13]. Still, pressures and genetic drift acting on particular mutations can foster distinct, functionally relevant differences between populations[14,15], even if they were nearly neutral in the original population. Slightly deleterious mutations can persist or be fixed[16–18], and epistatic interactions can cause neutral or beneficial variants in the original population to be harmful in the receiving population[7,9]. As these differences accumulate, incompatibilities may develop, contributing to reproductive barriers that prevent successful hybridization or limit its effect[19–21]. A hybridization event leads to two differentiated haplotypes producing hybrids and, therefore, exposes a plethora of variants to a new genomic landscape[22,23].

In baboons (genus Papio), hybridization between distinct populations is commonly observed. Still, our previous report[24] showed that genetic variants from foreign ancestry have to travel through the hybrid zone before reaching populations further from the hybrid zone[24–26], even in the face of extensive ongoing admixture. With its multiple species, the baboon species complex thus presents an ideal system for investigating negative selection mechanisms on admixture. Baboons have six distinct species, with the deepest split dated at least 1.3 million years ago between southern baboons (yellow, kinda, and chacma baboons; P. cynocephalus, P. kindae, and P. ursinus) and northern baboons (olive, hamadryas and guinea baboons; P. anubis, P. hamadryas and P. papio)[24,27]. A previous study by Vilgalys et al. on yellow and olive baboons found evidence of selection against admixture in a baboon hybrid zone in the Amboseli basin[26]. They found that gene-dense and low recombination regions had significantly less admixture than expected. In the hybrid zone, baboons with older admixture account for most of this correlation. In contrast, recent hybrids did not show this pattern, indicating that it takes at least 4-5 generations for introgressing chromosomes to break into segments of a size similar to the scale of recombination rate and gene density variation[28]. Negative selection on admixture may be due to deleterious variants introduced from an inbred population, maladaptation to local environments[2,29], and sexual incompatibilities reducing reproductive success[4,30,31]. Genetic incompatibility can be in the form of Bateson–Dobzhansky–Muller Incompatibility (BDMI), which may arise from negative epistatic interactions among genes. Selection against admixture from the rarer ancestry across swordfish populations has been reported as evidence of BDMI[3] by detecting negative selection on admixture[3,6,21,32,33].

Baboon species exhibit a striking discordance between autosomal and mitochondrial phylogenies, with a mitochondrial clade extending across East African olive, yellow, and hamadryas baboon ranges (dotted lines in Figure 1), even though olive and yellow baboons also each carry species-specific deeply diverged mitochondria in other parts of their species range. Our recent analysis[24] supported that this discordance arose when an ancient widespread hamadryas-like baboon was subsumed in yellow and olive baboon male-driven range expansions, displacing the nuclear genome of the resident population but not its mitochondrion. In this process, often referred to as nuclear swamping[34,35], this continuous displacement of original ancestry will result in the invading species going from minor to major ancestry. Any hybrid incompatibility would thus initially manifest as negative selection on the invading ancestry. As the invading ancestry reaches the majority, selection would change to favor the invading ancestry. The relative strengths of migration and negative selection on the minor parent ancestry thus determine which ancestry ends up fixed and could result in swamped populations retaining high-frequency original ancestry at incompatibility loci.

**Figure 1:**
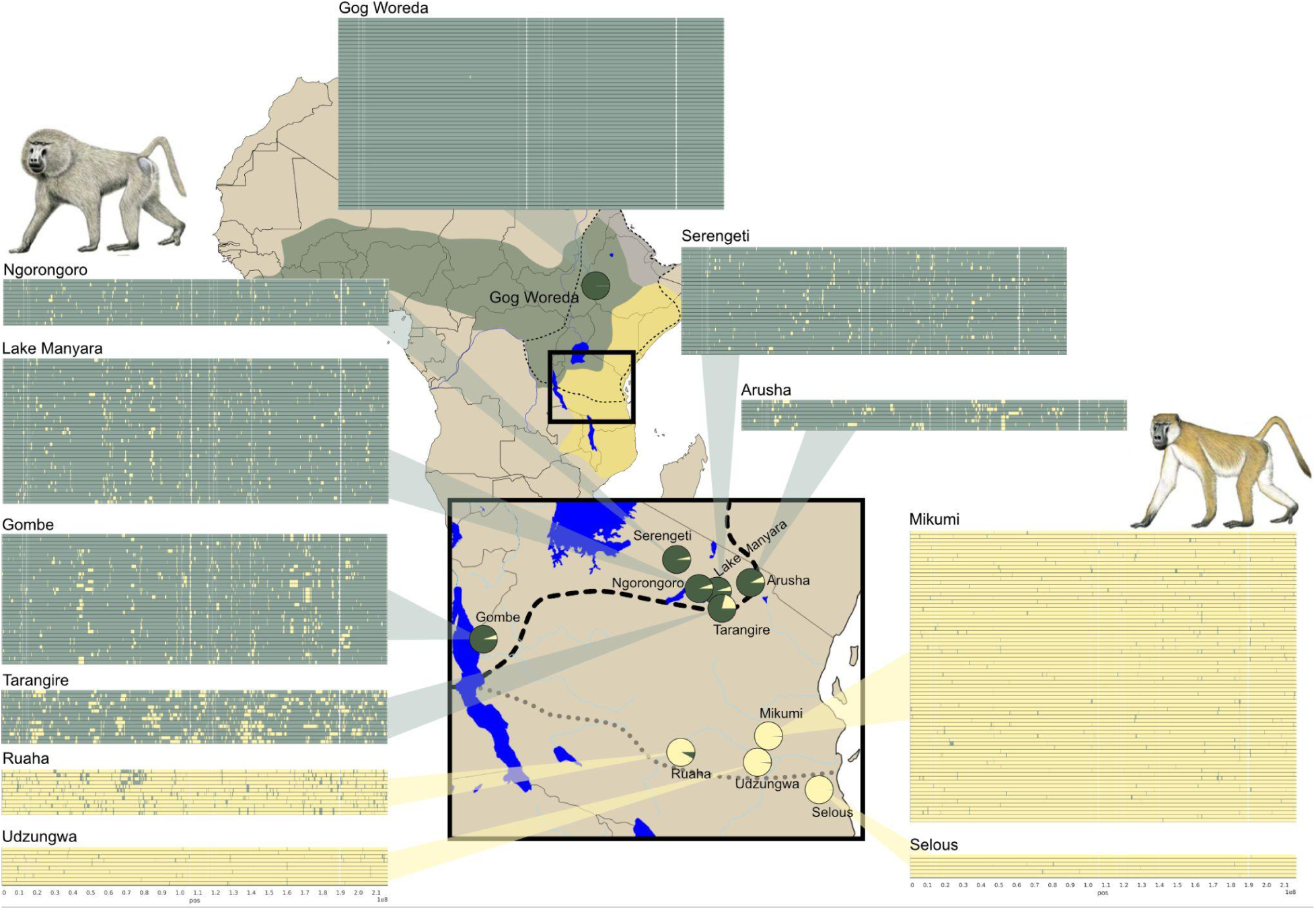
Overview of sampling locations and RFMix ancestry paintings of each individual’s phased chromosome 1 haplotypes. Olive and yellow colors represent northern and southern baboon ancestry. Pie charts at the sample locations show the MPA proportion of each population. The current olive/yellow baboon contact zone is shown with a dashed line. Dotted lines on Africa and insert maps show the extent of the mitochondrial G clade spanning the East African populations of yellow, olive, and hamadryas (hamadryas range shown in grey).

Haldane’s rule that genetic incompatibility first emerges in the hemizygous sex is supported by a century of observations. It suggests an outsize role of the mammalian X chromosomes in hybrid incompatibility[10,36–39]. The smaller effective population size (Ne) of the X chromosome results in faster divergence by genetic drift, a larger impact of recent changes in population size[40], and sensitivity to sex-biased dispersion and reproductive variance[38]. Dynamic co-amplification of multicopy ampliconic genes in *Mus musculus* and *Bos taurus* suggests that mammalian X and Y chromosomes participate in an arms race to control the X/Y ratio of fertile sperm[41–43]. This hypothesis predicts an accelerated evolution supported by multiple reports of positive selection on primate X chromosomes[44–47].

This study analyzes the relationship between admixture and linked selection across eleven sampling locations with varying distances from hybrid zones. The current species boundary represents a secondary contact between olive and yellow baboons[34,35,48,49]. Globetrotter was used to date the most recent admixture events to between 10 and 100 generations for the various populations[24,50,51] and found extensive migration between the olive baboon populations in Tanzania. The Amboseli population is similar to the Arusha and Tarangire populations, which all contain large amounts of minor parent ancestry. All three national parks are located near the boundary between olive and yellow baboons, with large admixture proportions in sampled individuals. Improved reference population size allows for better admixture inference, as parentage of rare variants, frequency of common variants, and species-specific haplotype combinations all are better resolved with more references. All individuals are 30X sequenced, allowing for higher accuracy and resolution methods. Sampling of a gradient of olive baboon populations shows how admixture patterns change as they penetrate deeper into the relatively pure olive baboon ranges. Unlike the previous studies[3,26], the X chromosome is also investigated, showing up to seven times stronger removal of minor parent ancestry in low recombination and low diversity regions.

We analyze a large dataset of phased high-coverage genome sequences first presented by[24,52]. There are 225 baboons sequenced at 30X across 19 localities. Whereas all 62 yellow baboons are sampled from the same general area in Eastern Tanzania, the 94 olive baboons are sampled from three general areas: Gog in Ethiopia, Gombe in Western Tanzania, Tarangire, Arusha, Ngorongoro, Lake Manyara, and Serengeti in Northern Tanzania. This study aims to elucidate the dynamics of selection against admixture in the baboon species complex, focusing on identifying genome-wide patterns and the role of the X chromosome. Chromosome X is known to have more admixture than the autosomes for both olive and yellow baboons[24], even though many other admixture cases exhibit decreased admixture on chromosome X[6,36,53].

## Results

### Minor Parent Ancestry declines with distance from the hybrid zone

We infer local ancestry along each high-coverage P. anubis (olive) and P.cynocephalus (yellow) baboon genome using RFMix[54]. The four other baboon species were used as reference panels. We assign olive baboon ancestry (“northern ancestry”) to genomic segments closer to P. hamadryas and P. papio (hamadryas and Guinea baboons) and yellow baboon ancestry (“southern ancestry”) to segments closer to P. kindae and P. ursinus (Kinda and chacma baboon). We consider minor parent ancestry (MPA) the rarer ancestry in each population and represent the MPA proportion along the genome as its mean in nonoverlapping, contiguous 100-kilobase windows. To assess the accuracy of RFMix, we used Haptools[55] to simulate admixture from the current standing variation of two of the sampled populations. The Pearson correlation between the simulated and admixture inferred by RFMix is 98.9%, with no bias towards major parent ancestry and high diversity regions (See Supplementary Simulation for details).

The MPA proportion of sampled populations differs widely (Figure 1, Supplementary Table 1). In yellow baboons, it drops from 8.0% in Ruaha, closest to the hybrid zone, to 0.36% in Selous, furthest away. In olive baboons, the MPA proportion drops from 20.9% in the Tarangire population, which borders the yellow range, to 3.4% in the Serengeti population, located 250 km into the olive range. This relationship with distance from the hybridization zone suggests that the introgressed sequence is removed by selection as it disperses by animal movement between populations close to the species contact zone[34,35,48,49], as this is not the original population ranges for yellow and olive baboons. Southern ancestry is virtually absent in the Ethiopian Gog baboon population, far from the hybrid zone (MPA proportion 0.065%), indicating little or no introgression from southern baboons into Gog olive baboons.

Recent incomplete lineage sorting (ILS) can contribute to estimates of MPA proportions[56,57]. However, ILS and old admixture events will mainly contribute MPA segments smaller than 0.05cM[58] that RFMix is not expected to detect[59]. Since ILS will affect all olive baboon populations equally, the absence of inferred southern MPA in the Gog population shows that RFMix inference is not biased by ILS and provides reliable estimates of recent admixture.

### Negative selection on minor parent ancestry

If the introgressed sequence carries variants under negative selection, we expect that more admixture is retained in high-recombination regions where neutral introgressed sequence segments are more readily unlinked from the deleterious variants that are eventually purged from the population[3,26]. Although we expect the rate of admixture removal to be highest in the first generations, where large numbers of negatively selected variants remain linked, we do not expect the correlation with recombination rate and MPA to be strong for very recent admixture where introgressed segment sizes remain orders of magnitude larger than that of fine-scale recombination rate and the 100kb scale at which we quantify MPA[28]. In these initial generations, removal will instead be much more indiscriminate due to the high linkage across the chromosomes[22,60]. We expect the strongest association between recombination rate and MPA as admixture disperses behind the hybrid zone and recombination reduces segment lengths to sizes relevant to medium-scale recombination rates and selection coefficients. However, the correlation will gradually weaken as the deleterious minor parent ancestry variants are removed and the MPA proportion is reduced.

To obtain fine-scale recombination rates, we created a genetic map with Pyrho [61,62] and SMC++ [63,64] using the 38 individuals from Mikumi. See Methods and Materials. The large number of samples from this population allows for a more accurate inference of the fine-scale recombination rate. However, the modest admixture will slightly bias our map toward lower rates in regions with higher MPA proportions. The recombination rate in 100kb windows has a mean and median of 0.0834 and 0.0668 centiMorgan (cM), scaled to a size equal to a previously published map for olive baboons of 2293 cM for the autosome[65].

The distribution of recombination in 100 kb windows has a rightward skew (see Supplementary section on outliers), and 0.5 % of 100kb windows have an estimated recombination rate above 0.408cM. When included, these outliers will have an outsized impact on the regressions. To restrict the analysis to a representative range of recombination rates, we removed the upper and lower 0.5 percentiles of recombination rate. We use variance-weighted linear regressions to account for heteroscedasticity (Supplementary Table 2 and Supplementary Figure 1) and find a significant association between the population mean MPA proportion in 100kb windows and the 100kb window recombination rate. This is true for all populations (p-values < 2e-5) except Tarangire, Arusha, and Lake Manyara (See Figure 2 and Table 1) and reveals negative selection on MPA in both olive and yellow baboon populations far from the hybrid zone. However, in line with expectations, the correlation is absent in Tarangire and Arusha, where admixture was most recently introduced, which is evident as longer MPA segments in the local ancestry paintings shown in Figure 1. Lake Manyara shows no significant correlation despite an MPA proportion smaller than in Ngorongoro and Ruaha.

**Figure 2:**
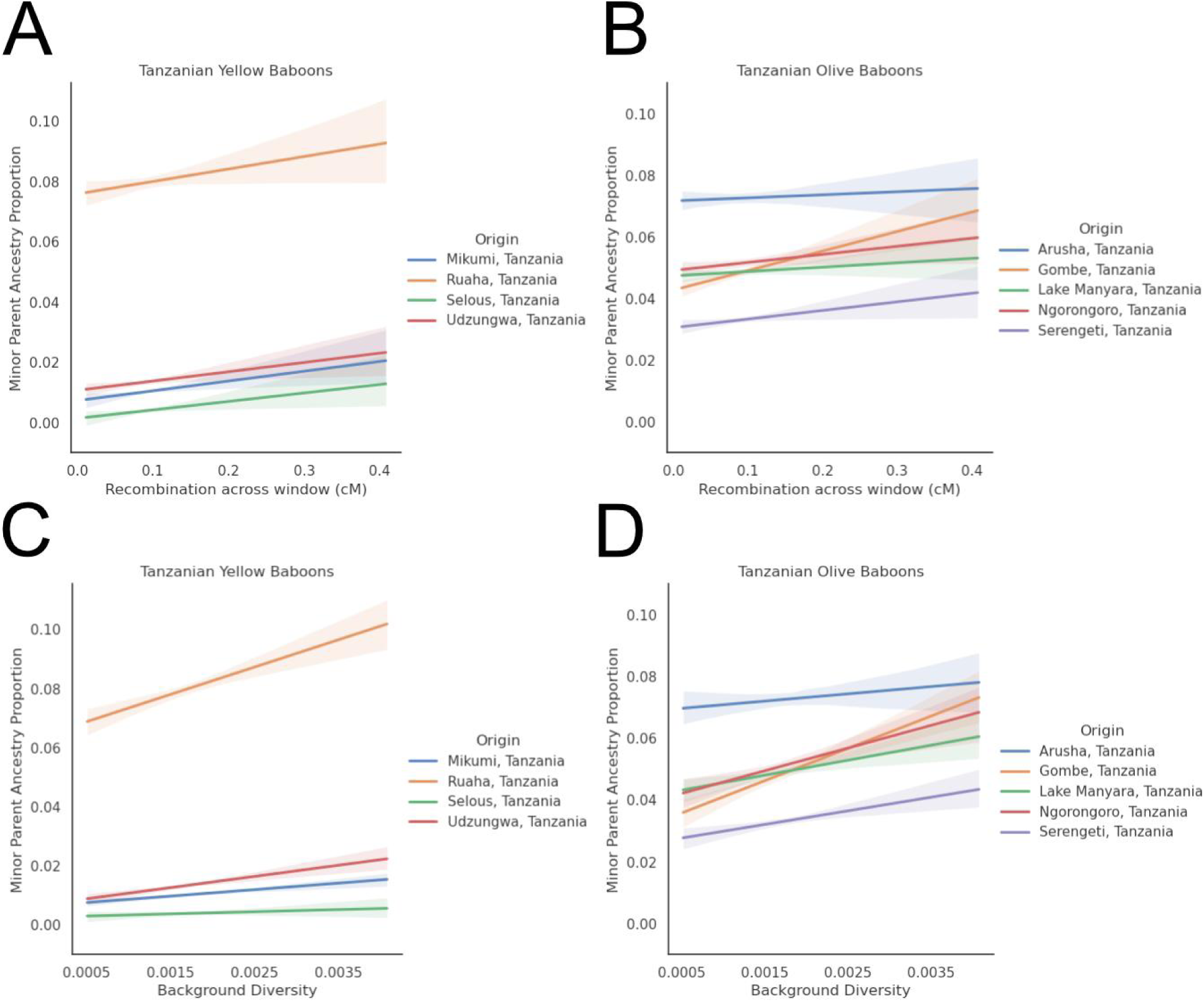
Weighted Linear Regressions for MPA proportion against recombination rate (A, B) and background diversity (C, D) Tarangire showed no correlation and is excluded for legibility.

Among yellow baboons, Ruaha, the most admixed population, has the largest regression slope (0.0414), significantly larger than for the Selous, Mikumi, and Udzungwa populations (p-value 0.00067) ranging from 0.0288 to 0.0325 (See Figure 2A, Supplementary Table 3). For an impression of the effect sizes, we ranked autosomal 100kb windows by their recombination rate and computed the mean MPA for windows in five equally sized bins ordered based on recombination rate (see Supplementary Figure 2 and Supplementary Table 4). Comparing mean MPA proportions for the 20% windows with the lowest recombination rates to the 20% with the highest rates shows a difference of 0.25% (0.896% to 1.15%), corresponding to a relative difference of 28%. The most admixed Ruaha population shows a similar absolute difference of 0.23 percentage points but a relative difference of only 3%. The lower relative differences for populations with higher MPA proportions are in line with the expectation that MPA is pruned most rapidly when the MPA proportion is high, as in Ruaha, and that it is continually pruned as the MPA proportion falls while becoming more discriminate, as shown by the larger relative difference in the Mikumi population.

The degree of admixture also varies among Tanzanian olive baboon populations (Figure 2B, Supplementary Table 1 and Supplementary Figure 2). The Gombe population has the steepest regression slope of 0.0633 (p-value 5.6e-19) for recombination rate and MPA proportion, and the Serengeti and Ngorongoro populations also have significant regression slopes (p-values 4.07e-07 and 0.00177). The populations in Tarangire and Arusha at the active contact zone do, as expected, not show a significant relationship between recombination rate and MPA. Arusha has the largest absolute and relative difference between low and high-recombination windows (6.82% in the lowest 20% to 7.22% in the highest 20%, a 6% increase). Ngorongoro has 4.92% MPA in the lowest 20% to 5.16% MPA in the highest 20%, a 5% increase. See Supplementary Table 4 for all quintile differences. We find no significant correlation between recombination rate and MPA proportion in Lake Manyara (p-value 0.0306).

As a proxy for linked selection, the recombination rate does not account for variation in the density of sites under negative or positive selection. A more direct quantification of linked selection is relative genetic diversity, measured as π, the mean pairwise differences among haplotypes in the population. Although recent bottlenecks may exacerbate variation in diversity along the chromosomes[66], the relative degree of diversity is primarily shaped by linked selection[60,67–70] and variation in mutation rate. However, as admixture itself contributes to 100kb mean diversity, we must obtain independent estimates of π from “background” populations, which do not share historical admixture events with the studied population. In the absence of admixture in the background populations, this estimate of π quantifies linked selection as a proxy for the functional importance of each 100kb window. If independent admixture contributes to background population diversity, its distribution across 100kb windows will additionally reflect the strength of negative selection on MPA; i.e., diversity measured in admixed background populations is expected to correlate more strongly with MPA proportion in the study population if they also are admixed[24]. We calculated the average diversity across Guinea, Hamadryas, Kinda, and Chacma baboon reference populations in 100kb windows and will refer to this as “background diversity” below. (Supplementary Figure 3). Background diversity strongly correlates with the recombination rate (Pearson correlation: 0.602, p-value: 0.0, Supplementary Figure 4).

Among yellow baboons, the Ruaha population again shows the highest regression slope (9.25, p-value 1.91e-19), significantly higher than other yellow populations (p-value: 1.01e-10), while the Mikumi (slope 2.19, p-value 1.21e-16) and Udzungwa populations also have significant regression slopes and slope 3.79, p-value 9.51e-18) (Supplementary Table 5). The mean MPA proportion in the bottom and top 20% windows ranked by background diversity (see Supplementary Figure 5 and Supplementary Table 6) represents an absolute difference of 16% (7.68% to 8.92%). The Mikumi and Udzungwa populations also show significant correlations (p-value 1.21e-16 and 9.51e-18), Udzungwa showing the largest relative difference, 44%, between the bottom and top 20% windows (1.22% to 1.76%). Further from the hybrid zone, the Selous population shows no significant correlation after Bonferroni correction to account for testing multiple populations (see Figure 2C).

Among the olive baboon populations, the correlation between MPA proportion and background diversity is significantly stronger in populations away from the hybrid zone. Gombe (p-value 4.35e-34) and Ngorongoro (p-value 4.03e-13) show the strongest correlation between background diversity and MPA (see Figure 2D). The Gombe population has the largest absolute and relative span in MPA proportions (4.46 % in the lowest 20% to 5.74% in the highest 20%, a 29% increase). The Ngorongoro population has the second largest absolute difference (5.14% in the lowest 20% to 5.65% in the highest 20%, a 10% increase). The Tarangire and Arusha populations closest to the contact zone show no significant correlation between background diversity and MPA.

A way to quantify whether there is a relationship between admixture level and selection against admixture is to ascertain whether background diversity on individual chromosomes correlates with their mean MPA proportions. (See Supplementary Figure 6). In general, the MPA is not randomly distributed but instead shows enrichment of MPA in the half of the chromosome with high background diversity, and this relationship is stronger on chromosomes with low relative admixture.

To compare the predictive strengths of background diversity and recombination, we z-normalized each one and used a generalized linear model to explain MPA proportion as a function of both recombination rate and background diversity (see Supplementary Table 7 for all coefficients and p-values). Of the yellow baboon populations, Mikumi, Udzungwa, and Ruaha all had a significant association with background diversity in predicting MPA proportion (p-values 0.00181, 3.4e-10, and 1.4e-15). For the olive baboon populations, Ngorongoro, Gombe, Lake Manyara, and Serengeti also had significant regression slopes (p-values 2.48e-06, 4.45e-26, 9.22e-10, and 3.43e-12) for background diversity and MPA. The Ruaha, Gombe, Lake Manyara, and Serengeti baboon populations had a significant negative association with recombination for MPA prediction (p-values 0.000113, 2.43e-07, 2.78e-07 and 6.86e-07) while maintaining a larger positive slope based on background diversity. As an example of the different regression slopes, the slope of the Ruaha population, when using only recombination, was 0.0414. When using only background diversity, it was 9.25, whilst the regression using both had slopes of -0.0559 and 10.5, respectively. That is, while the recombination rate is positively correlated with the MPA proportion, its correlation becomes negative when modeled together with background diversity.

### Stronger selection against admixture on the X chromosome

In all studied populations, the diversity on chromosome X relative to autosomes is significantly lower than the ¾ expected from hemizygosity[71]. This is partly explained by recent bottlenecks experienced by many populations (See Supplementary Figure 7), most severely in Northern baboons, which more strongly affect chromosome X diversity. However, even after adjusting for the effect of historical population sizes and the lower mutation rate on chromosome X[72], the X/autosome ratio remains significantly below ¾ (See Supplementary Table 8) (Mann-Whitney U test, p-values from 1.78e-26 to 2.17e-272). The X/autosome ratios range from 0.346 to 0.521, corresponding to a diversity reduction of 11.8% to 36.6% beyond expectation when adjusting for population size history and mutation rate. All baboons, except hamadryas, exhibit male-biased dispersion and higher reproductive variance in males[34,38,73–77], which should contribute to a higher X/autosome ratio. This leaves stronger linked selection on the X chromosome as the only explanation for the lower ratio. See Supplementary Figure 8 for the distribution of background diversity on chromosome X.

All olive baboon populations, except Tarangire and Arusha, show significant correlations of MPA proportion with both background diversity (Figure 3B and Supplementary Table 9) and recombination rate (Supplementary Table 10) on chromosome X. The Ngorongoro population has the strongest association between background diversity and MPA proportion (slope 81.1, p-value 8.19e-09). Here, the mean MPA proportion goes from 0.47% to 7.27% between the bottom and top 20% 100kb windows ranked by background diversity, a 14.4-fold difference. The similarly strong association in the Serengeti olive baboons (slope 68, p-value 2.47e-08) shows a span of 0.70% in the lowest quintile to 6.37% in the highest quintile in MPA proportion, an 8.04 fold difference. Among the yellow baboon populations (Fig 3A, Table 3), only the Ruaha population shows a significant association between background diversity and MPA proportion (slope 156, p-val 1.79e-16). Here, the 100kb windows with a background diversity in the top and bottom quintiles are 7.94% and 20.1%, a 2.5-fold difference (See Supplementary Figure 9 and Supplementary Table 11). The overall admixture in low-diversity regions is similar amongst all four yellow baboon populations. Still, the Ruaha population has much more admixture in high-diversity regions than the other yellow baboons. After Bonferroni correction, the Selous, Mikumi, and Udzungwa populations do not show significant correlations to either MPA proportion or recombination rate (See Supplementary Table 9 and 10).

**Fig 3:**
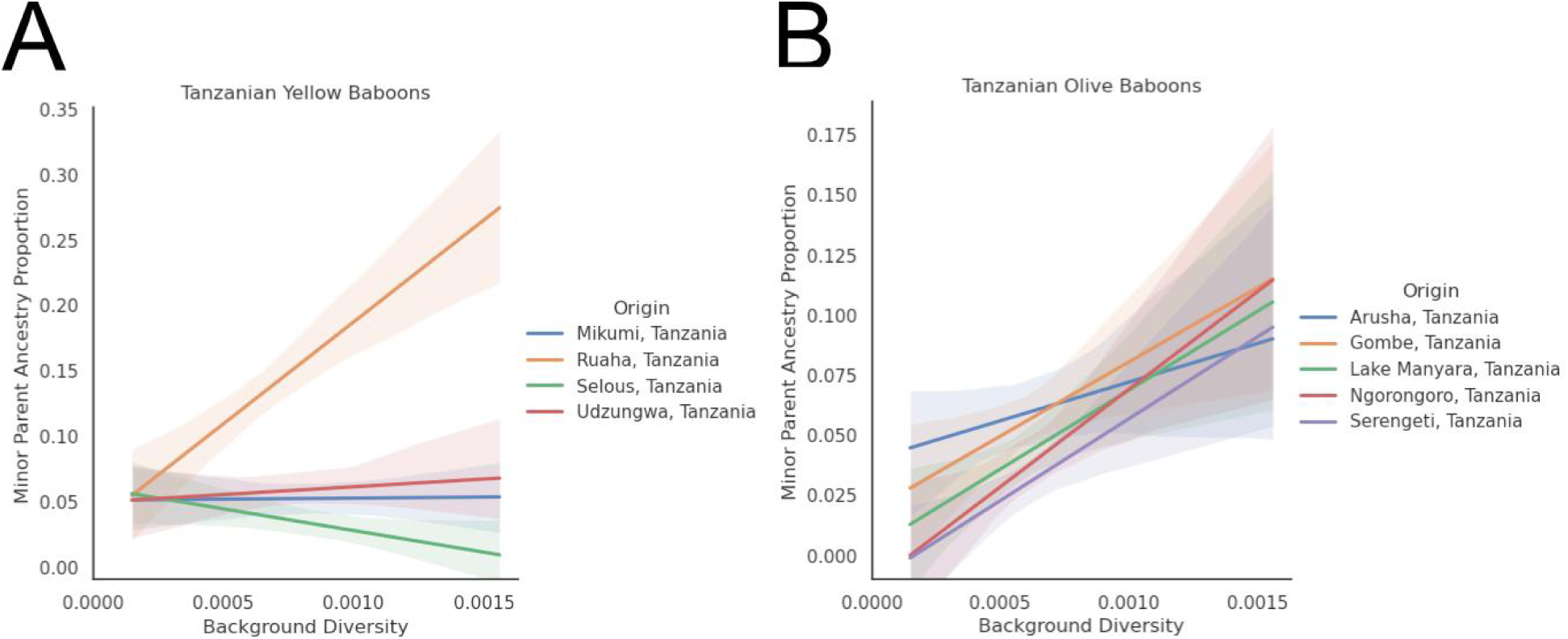
A-B) Weighted Linear Regression of MPA proportion against background diversity on the X chromosome.

To address whether the correlation between background diversity and MPA proportion is stronger on the X chromosome than on autosomes, we Z-score normalized background diversity and performed weighted linear regression MPA proportion against normalized diversity, chromosome type, and their interaction, keeping the variance weights the same as in the previous models. A linear model with an interaction term can determine whether there is a significant difference between the autosomal regression’s slopes and chromosome X’s.

In olive baboons, the slopes for the Ngorongoro, Lake Manyara, Serengeti, and Gombe populations are significantly higher or the X chromosome (p-values 6.29e-06, 2.53e-06, 1.42e-09, 0.00235). The effect is strongest in Ngorongoro and Serengeti, where slopes are 5.53 and 7.13 times steeper on the X chromosome. Tarangire and Arusha (Figure 4) do not have a significant difference between chromosome X and the autosomes, indicating again that the hybrid zone selection is too wide-reaching to segregate in 100kb windows. The Ruaha yellow population (Figure 4) also shows a significantly larger slope on chromosome X, 7.4 times steeper than for autosomes (p-val 1.42e-14). Selous has a negative interaction (p-val 2.72e-08) only due to the slightly negative autosomal slope. Mikumi and Udzungwa do not have any significant interaction (See Supplementary Table 12).

**Figure 4:**
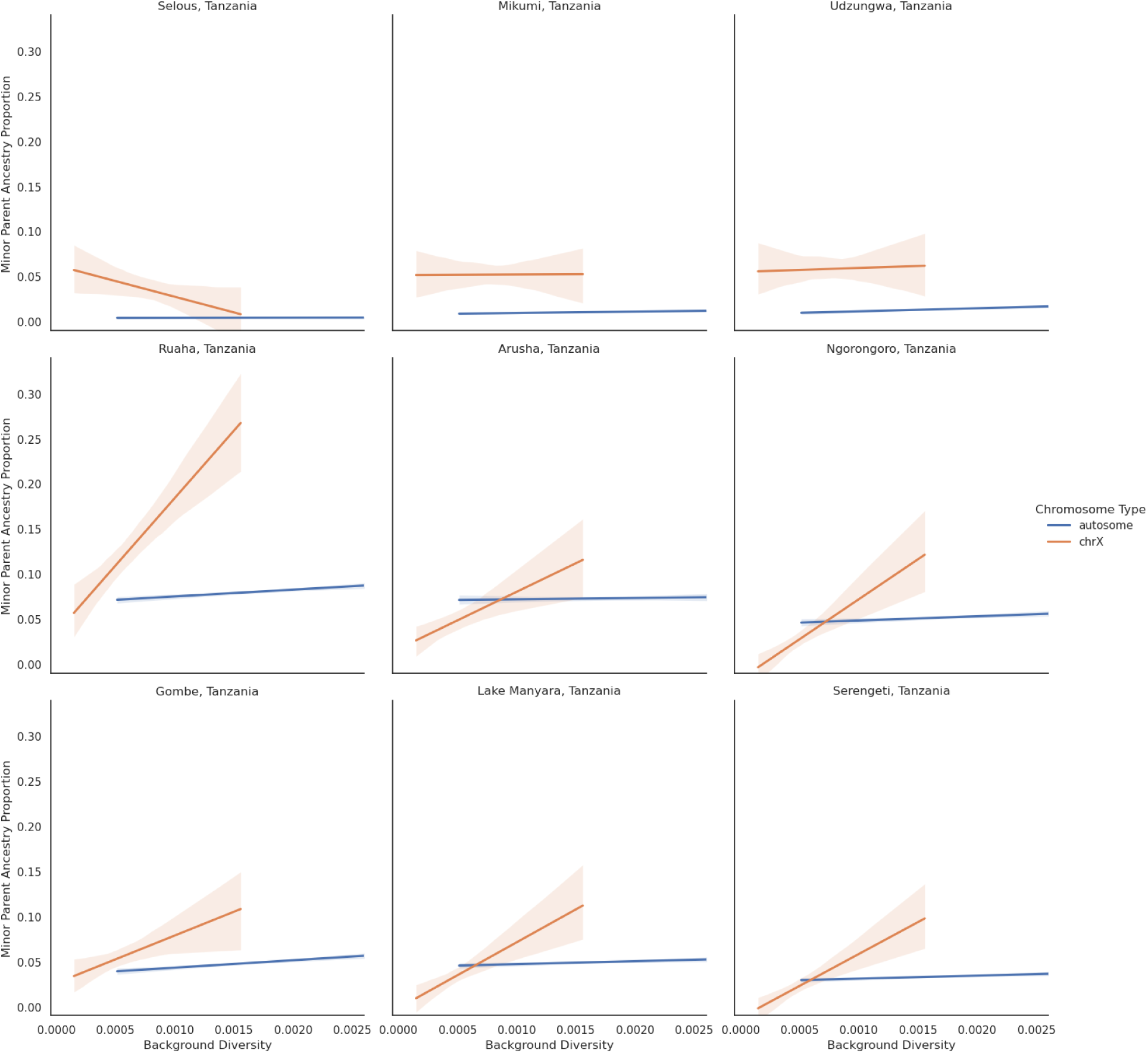
A truncated, unnormalized plot of background diversity and MPA on the autosomes and chromosome X for all Tanzanian baboon populations except Tarangire.

In all yellow baboon populations, the MPA proportion is significantly higher on chromosome X than on the autosomes (Figure 5, Mann-Whitney U-test p-values below 8.06e-05, see Supplementary Table 13). The olive baboon populations show no significant difference (See Figure 5). Chromosome X MPA proportions vary between 25.0 % in Tarangire and 4.2 % in Serengeti (see Supplementary Table 1). MPA proportions in the yellow baboon populations vary between 12.9 in Ruaha and 4.5% in Selous. In autosomes, the corresponding variation spans 8.0 % to 0.36 %. Supplementary Figure 10 for the distribution of MPA proportions across individual chromosome haplotypes, stratified by chromosome number, and Figure 6 for visualization of RFMix inference for chromosomes eight and X.

**Figure 5:**
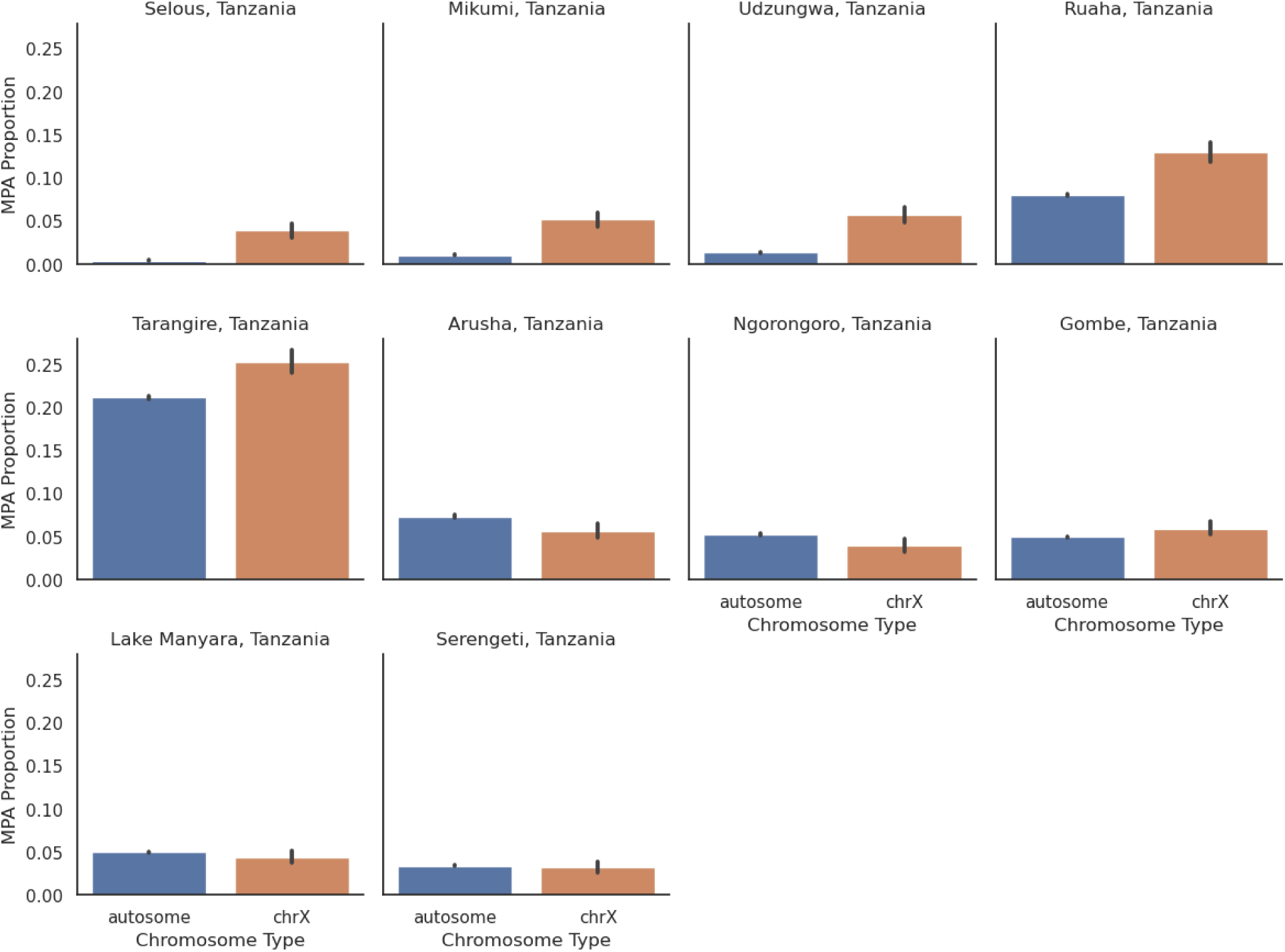
Barplot of autosomal and X-linked Minor Parent Ancestry Percentage for yellow and olive baboons in Tanzania.

**Figure 6:**
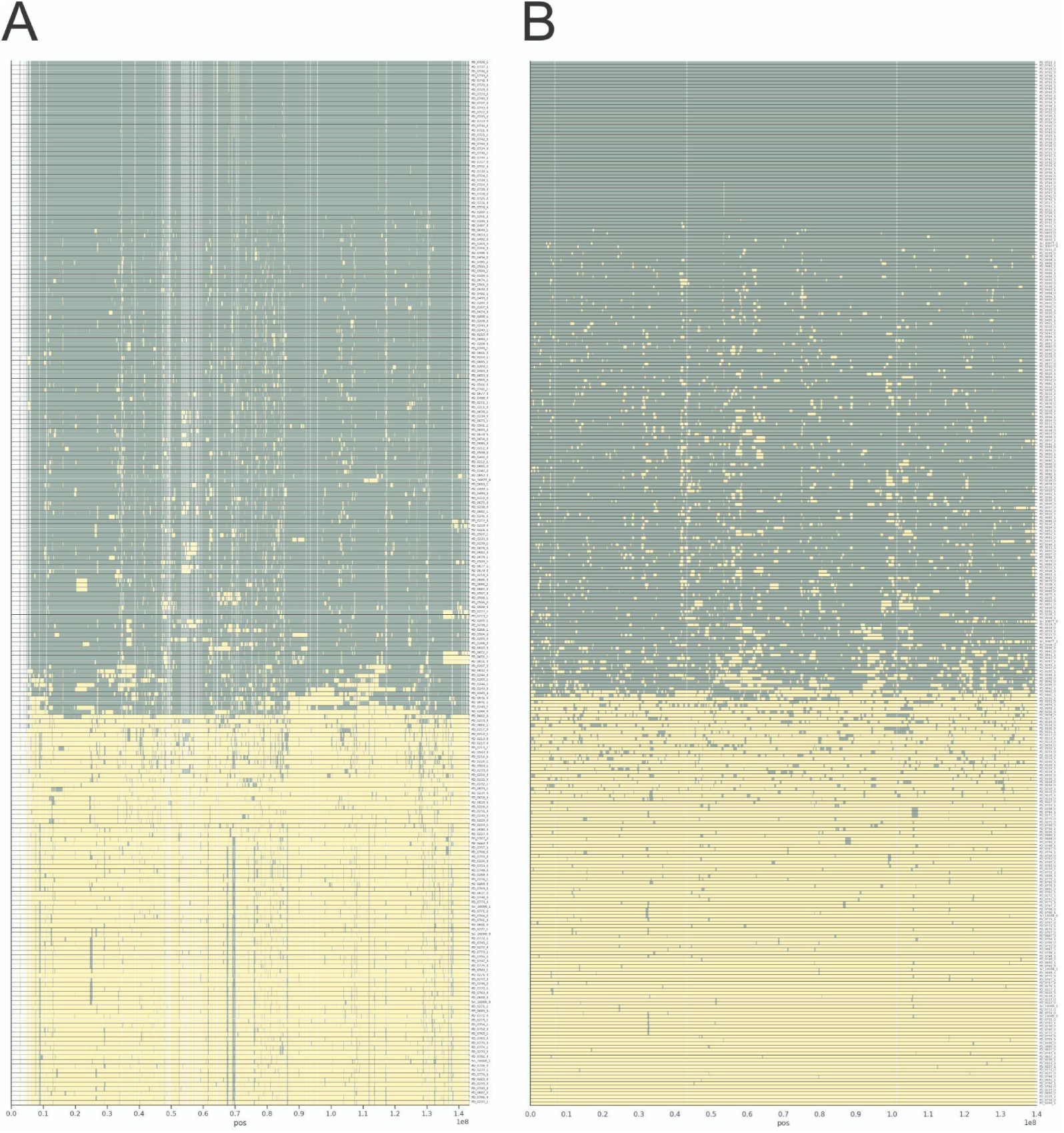
**A)** RFMix ancestry paintings of each individual’s phased chromosome X haplotypes. Olive and yellow colors represent northern and southern baboon ancestry. The haplotypes are clustered on similarity using UPGMA and otherwise sorted by sampling location. Windows with less than 75 % callability are filtered out. **B)** Same as A), but here the similar sized chromosome 8 for comparison. Note that there are fewer haplotypes for Chromosome X due to males only carrying one.

The higher MPA proportions on the X chromosome of many populations seem to contradict our observation of stronger negative selection on admixture on the X chromosome. On the X chromosome, the distribution of MPA proportions is skewed toward higher frequencies (See Supplementary Figure 11), particularly in the yellow baboons, where it is also evident as vertical patterns in the local ancestry painting (Figure 6). To investigate if high-frequency MPA is responsible for the stronger correlations on chromosome X, we repeated the analysis after masking 100kb windows with MPA proportions above 25% (See Supplementary Figure 12). Individual X and autosome regressions remain significant, but the association on X is strengthened in yellow baboons and has the opposite effect in olive baboons (Supplementary Table 14): The slopes for chromosome X are now higher in Udzungwa and Mikumi yellow baboons and are now significantly higher than those for autosomes (p-values 7.26e-15 and 2.35e-05), whereas Lake Manyara and Serengeti olive baboons no longer show higher slopes on chromosome X relative to the autosomes. This observation suggests that any selection on high-frequency MPA in yellow baboons is not represented by background diversity.

To investigate if high-frequency MPA could explained as recent adaptive introgressions, we used Relate to scan for recent selective sweeps. The sample sizes of each population are not sufficient for this analysis. However, an analysis of the pooled Tanzanian olive baboon populations yielded significant evidence of positive selection across the autosome but none on the X chromosome (See Supplementary Figure 13, Supplementary Section Positive Selection Scan)

### Hamadryas MPA in Ethiopian olive baboons reveals strong selection on hybrid ancestry

The observation that high-frequency MPA is most common on X chromosomes of the yellow populations that represent nuclear swamping of a resident Northern lineage suggests that it may be remnants of the original population retained at high-frequency by negative selection on incoming yellow baboon MPA. To explore this scenario further, we also analyzed the separate hamadryas MPA component of the Gog olive baboon population in Ethiopia. Olive and hamadryas baboons belong to the northern clade of baboon species and are thus more similar than olive and yellow baboons. Their divergence is comparable to that of anatomically modern humans and Neanderthals[27,78], The mitochondria of the Gog olive baboon population are more similar to those of hamadryas baboons than to the mitochondria of other sampled olive populations, including the Tanzanian olive baboons, which also carry a hamadryas-like mitochondria[24]. This has been explained by nuclear swamping, where an invasion of primarily olive males displaced autosomal but not mitochondrial ancestry of a resident hamadryas-like population.

We ran RFMix to infer hamadryas-like ancestry along chromosomes of the Gog individuals. We used Tanzanian olive and Hamadryas baboons as reference panels. Minor Parent Ancestry (MPA) is calculated as the proportion of Hamadryas ancestry in 100 kilobase windows after filtering areas with less than 75% callable bases (See Materials and Methods). Gog olive baboons have more admixture from hamadryas than any Tanzanian population from southern baboons and a larger proportion of high-frequency MPA than the other investigated populations (Fig 7D, Supplementary Fig 14).

**Fig 7:**
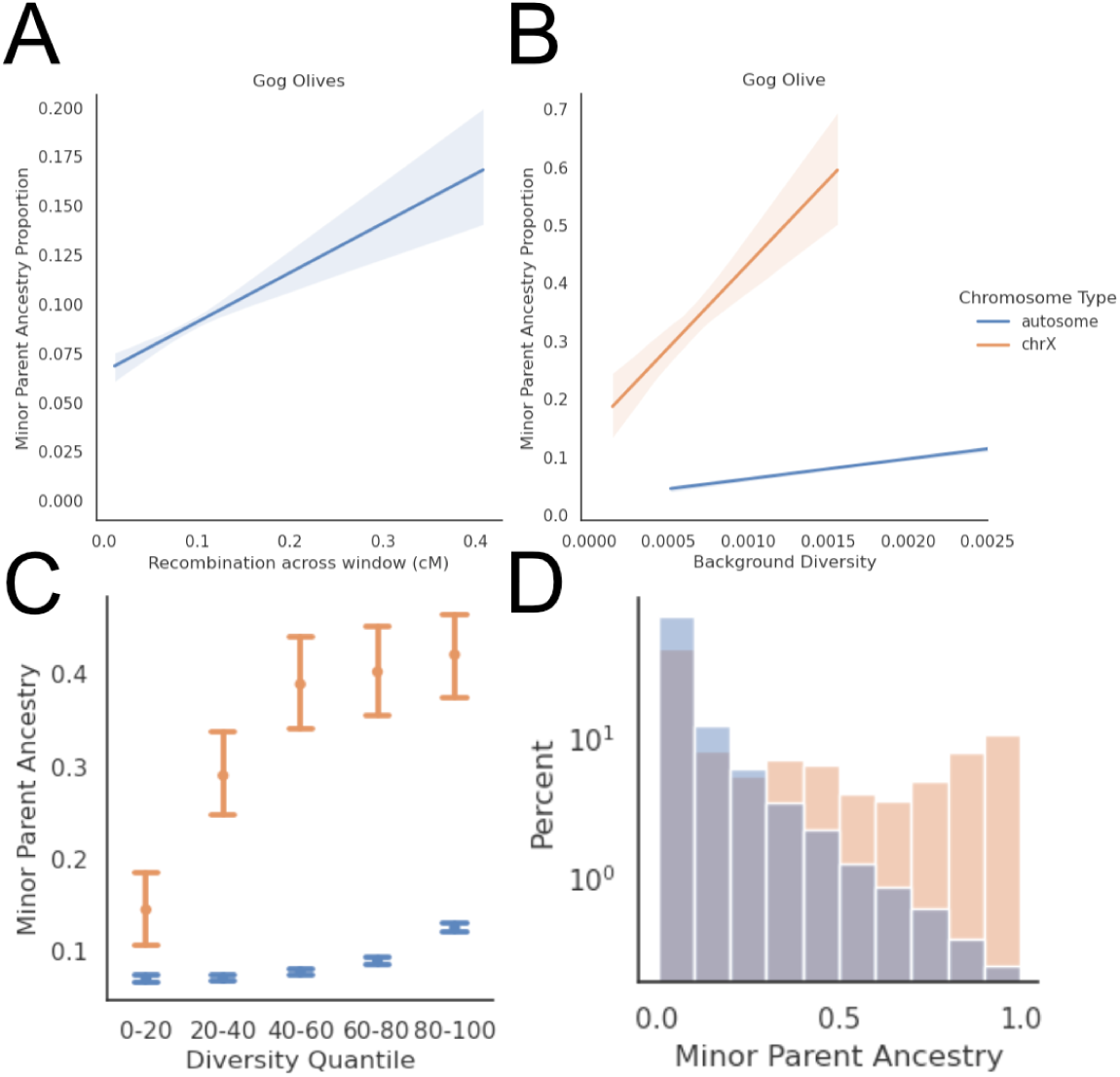
**A)** Weighted Linear Regressions of MPA proportion along Gog olive X chromosomes against recombination rate.. **B)** Weighted Linear Regressions of MPA proportion along autosomes and the X chromosome against background diversity.**C)** MPA proportions for autosomes and the X chromosome in bins of background diversity **D)** Histogram of log-scaled MPA proportion across autosomes and the X chromosome.

On the autosomes, Gog olive baboons have the steepest slope of all investigated baboon populations against both recombination (Figure 7A, slope 0.251, p-value 1.03e-76) and background diversity (Figure 7B, slope 38.8, p-value 1.58e-132) (Supplementary Table 15), as well as the largest absolute difference between the lowest 20% background diversity quintile and the highest (7.05% in the lowest 20% to 12.6% in the highest 20%, a 78% increase). With a regression using both recombination rate and background diversity, it is only background diversity which has a significant association (p-values 0.393 and 1.41e-65).

Despite a large MPA proportion (Fig 7C and D), the X chromosome of Gog olive baboons also shows the steepest slope against the MPA proportion of all investigated populations, slope 214, p-value 7.18e-10), significantly higher than that for autosomes (p-val 8.88e-06). After normalization, the X chromosome regression is 54% steeper than the autosomal one. See Supplementary Table 15 for all regressions. Gog olive baboons also show the largest absolute difference between the 20 % low diversity quantile chromosome X (14.5% to 42.1%, a 190% increase). These observations are qualitatively similar to those for Yellow and olive admixture but are greatly amplified: The evidence of negative selection is strongest here despite an extreme MPA proportion of 33.1%, suggesting a selection regime different from autosomes where the proportion is only 8.8%. These observations are all predictions of nuclear swamping and point to strong selection retaining original ancestry on the X chromosome.

## Discussion

Reproductive isolation leads to divergence. If isolated for long enough, the populations will become different species. Baboons are deeply diverged, but they still interbreed when they meet. A previous approach to investigate possible selection against admixture is to see whether recombination rate can predict levels of minor parent ancestry, as recombination rate determines how quickly neutral variants can be decoupled from deleterious variants and, therefore, survive in the new selective landscape it finds itself in when admixture occurs. In three distinct admixture cases, with olive, yellow, and hamadryas baboon minor parent ancestry, recombination rate can be used as a significant predictor of admixture degree across the genome.

This association between recombination rate and MPA proportion is found across multiple populations and arises when moving away from the hybrid zone. This association can also decay if not enough admixture is present. Low degrees of observed admixture correlate with old admixture events in this dataset, and rare bouts of positive selection penetrating deeper into species ranges will be observed as a large proportion of admixture when MPA proportion is low. Selection against admixture is a complex process, and the resulting admixture patterns depend strongly on how admixture occurs, such as whether it is a sudden pulse or protracted, how old the admixture fragments are, and how diverged the populations are. The admixing haplotype will be divergent from the haplotypes of the receiving population and contain more variants that may be under selection. This density of selected variants might cause a full purge of admixture, which affects local diversity less than weaker background selection[70,79]. Therefore, the strongest association between recombination rate and MPA proportion is expected in population combinations in which there will be persistent selection against admixture without very large selective effects, as highly deleterious alleles will cause extremely widespread purging of admixture which cannot be detected with these statistics.

We also show that the diversity patterns in unrelated populations are a better predictor across all populations except Selous yellow baboons. Recombination rate, while useful, only has little information on the density of selected variants, while diversity patterns depict how frequently linked selection depresses variation. In both cases, the local mutation rate is not accounted for, which varies based on the local context in the genome[72,80–82], and possibly would improve predictions even further if paired with background diversity and recombination. When performing a regression with both background diversity and recombination, many populations had a significant positive association with background diversity. In contrast, the recombination rate had either no significant association or a negative one, except for Selous. This indicates that when adjusting for diversity patterns, the recombination rate overestimates MPA in high-recombination rate regions or underestimates MPA in low-recombination rate regions. This is probably due to the three factors influencing linked selection: mutations, density of selected variants, and recombination. Mutations cause linked selection, if not neutral, mediated by recombination[60,67]. However, there is also another pathway, which is that recombination itself causes mutations[83–85]. The recombination rate is, therefore, correlated with the mutation rate, which is variable across the genome. Being able to adjust for recombination, therefore, shows a negative correlation, as it removes one of the confounders that can cause high diversity while linked selection still may act in that region.

The effect of selection against admixture requires a balance between the presence of admixture and the time for selection to act to remove it. If the admixture event is too old, the effect of drift and positive selection of specific variants can overpower the effect of negative selection against admixture. In yellow baboons, Selous, with a low degree of admixture, shows no association with background diversity, while Ruaha, with the highest MPA proportion, shows the strongest relationship. Meanwhile, Tarangire and Arusha, the two most admixed olive populations, show no correlation between MPA and background diversity after multiple testing correction. Depending on the predictor used, the Ngorongoro and Gombe populations, with intermediate levels of MPA, have the strongest correlation among olive baboons. The Gombe olive population admixes with a markedly different population than the other olive baboons, as it is near a more Kinda-like population in Mahale and Katavi[24] with a substantially different ancestry than the yellow baboons in eastern Tanzania. For chromosome X, the populations with the most significant associations are closer to the hybrid zone than is the case for the autosomes, with Ngorongoro showing the largest difference between autosomes and chromosome X.

An important difference when considering selection against admixture is whether an admixture event can best be described as a minority admixture event, which always is the minor parent ancestry after the hybridization, or a nuclear swamping event, wherein repeated dispersal into a receiving population eventually changes the ancestry into majority migrant ancestry[34]. As shown in all three admixture cases, there is more admixture on chromosome X, indicating that nuclear swamping is in play in all three cases, as it primarily is males that migrate in all baboon species except hamadryas[34,77].

In most cases, chromosome X has a stronger correlation with background diversity, even though chromosome X has more admixture than the autosome in all populations except Ngorongoro. Some of this difference might be because olive and yellow baboons exhibit male-biased dispersal and, therefore, might take over habitats through male-biased admixture. Chromosome X in baboons is also strongly depressed in diversity compared to the expectation, which can only occur if there is much more selection and/or biased mating patterns in baboons, in which females are more reproductively variable than males, which is unlikely[34,73,74,86]. The opposite is generally observed, with a few males ranking highly in baboon troops and mating more often than the rest.

The most striking difference between chromosome X and the autosomes is in Gog olive baboons. These baboons carry mitochondria more similar to hamadryas baboons than other olive baboons, indicating that hamadryas baboons previously inhabited this area or that hamadryas females are migrating into this location. This might partly explain why chromosome X is enriched for admixture, but it can’t be the only explanation due to the large difference in admixture level and selection against admixture. Stronger selective pressure on chromosome X purging minor parent ancestry may explain this pattern. First, as nuclear swamping progresses, chromosome X would maintain higher levels of the original ancestry until autosomal ancestry became the majority admixing side when selection against admixture would flip to removing the original ancestry. Therefore, selection against minor parent ancestry on chromosome X would be delayed compared to the autosomes. If that is the case, it can only be explained by BDMI, not by ecological adaptions or sexual selection. This pattern of chromosome X incompatibility is strongest in Gog olive baboons, but it is also found in other high-admixture cases, such as the Ruaha, Ngorongoro, and Gombe populations.

To see a significant correlation between diversity and admixture, selection against admixture has to be in an intermediate state, with strong enough selection to distort admixture patterns but weak enough not to be purged in the first generations after admixture. Therefore, the correlations can be stronger in an admixture case where there is less selection against admixture, as regions with very large selection coefficients are rarer, and smaller selection coefficients affect neighboring areas more than variants removed immediately[60,67,87].

To conclude, baboon populations in four different localities, with yellow baboons in southern Tanzania and olive baboons in western and northern Tanzania as well as Ethiopia, all show a significant association between admixture and recombination/diversity of background populations, indicating extensive selection against admixture no matter the minor/major parent combination. Leveraging both recombination rate and background diversity shows a negative correlation with recombination rate in some populations, indicating that background diversity association would be further improved if the mutational landscape could be included. In addition, the X chromosome both has more inferred admixture and a much stronger selection process against it, supporting the hypothesis that faster X chromosome evolution makes it a prime target for incompatibilities, as well as the hemizygous nature causing increased removal of deleterious alleles in the minor parent ancestry. We also show that the decreased diversity on chromosome X can only be because of increased negative selection, and selective sweeps inferred by Relate[88] are enriched in areas with low or no admixture.

## Materials and Methods

A mask was created based on the depth of coverage per individual. For each individual, the most common read depth (mode) was calculated using bcftools (see Sup Fig 15 for distribution, Supplementary Section Commands). Each base was set as passing if the read depth was between one-third (min_cov) or twice (max_cov) the mode for that chromosome. Sites were filtered using bcftools, removing sites that were uncalled, had a heterozygous call and alternate allele depth less than 3, coverage less than a third of the mean coverage or more than twice, or a GQ score less than or equal to 30. A species-wide mask was created by calling a base as passing if 95 % or more of samples had a passing state with bedtools multiIntersectBed. For chromosome X, the mask was created using only the 98 females.

Diversity was calculated using scikit-allels windowed_diversity function with a window size of 100kb. All windows in the analysis are 100kb, starting from base 1 of the reference genome unless stated otherwise. Windows were removed from the analysis if less than 75% of the window was set as passing in the filter step.

Historical effective population size for every population was inferred with SMC++. The autosome mutation rate was set at 0.57e-08 per base and 4.5 times more mutations in males[72], while chromosome X was set at 0.45e-08. This mutation rate was calculated based on the higher mutation rate in males corresponding to (4.5/3+2/3)/(4.5/2+2/4), the mutation rate in chromosome X divided by the autosome mutation rate. Generation time was set at 11 years[72]. The Autosomal and chromosome X SMC++ run was run with a piecewise spline and ten expectation maximization iterations. Chromosome X Ne was inferred only using females.

A fine-scale recombination map was inferred using Pyrho and the Mikumi Yellow population, as it had the largest sample size. The Pyrho workflow consists of three steps: making a lookup table, inferring optimal hyperparameters, and then inferring the fine-scale recombination. Pyrho (maketable) was run with standard options except for the approximate setting for computational feasibility. The autosomal and chromosome X population size histories for Mikumi yellow baboons were used. Pyrho (hyperparam) was run with the following possible block penalties and window sizes: 10,25,50,100 and 10,25,50,100.

Then, Pyrho (optimize) was run with optimal hyperparameters as measured by the L2 norm for each chromosome. Lastly, the autosome-wide recombination rate was scaled to be equal to the estimate of 2293cM published by [65], as Pyrho is optimized to infer fine-scale recombination and not total genetic distance. The scaling factor used was 1.3, as the total autosome-wide recombination rate inferred from pyrho was 1761. The assembly used for this estimate is Panubis1.0, while the one used in [24] is panu3. The recombination rate per window was calculated based on the Pyrho genetic map, based on the genetic distance between the first and last base in the window, with interpolation performed assuming an average recombination rate between SNPs.

Local Ancestry Inference was done using RFMix version 2, removing all alleles with a Minor Allele Frequency of less than 1 % for computational efficiency. RFMix was run with the Pyrho genetic map and an assumed admixture date of 100 generations ago.

Relate was run with a mutation rate of 0.57e-8 and an effective initial population size of 50000 for the autosomes, and chromosome X with a mutational rate of 0.46e-8 and an effective population size of 25000, in both cases with a generation time of 11 generations. The callability mask is the same as previously detailed. Otherwise, standard options were used for the Relate workflow, with the convert from VCF and prepare input script used to transform the VCF into haps/sample format, followed by Relate, EstimatePopulationSize, and then DetectSelection.

We used weighted regressions to account for heteroscedasticity, as identified with the Breusch-Pagan test[89]. We used the statsmodels python package to run the regression models

## Author Contributions

Conceptualization: All authors. Data curation: E.F.S. Formal analysis: E.F.S. and K.M. Investigation: E.F.S., and K.M. Methodology: E.F.S. Supervision: G.H., and K.M. Visualization: E.F.S., and K.M. Writing – original draft: E.F.S., and K.M. Writing – review & editing: All authors.

## Acknowledgements

We thank Mikkel Heide Schierup and Dietmar Zinner for feedback on the manuscript, and Jeffrey Rogers, Christian Roos and Clifford J. Jolly for insightful discussion.

## References

1. Marques DA, Meier JI, Seehausen O. A Combinatorial View on Speciation and Adaptive Radiation. Trends Ecol Evol. 2019;34:531–44.

2. Worsham MLD, Julius EP, Nice CC, Diaz PH, Huffman DG. Geographic isolation facilitates the evolution of reproductive isolation and morphological divergence. Ecol Evol. 2017;7:10278–88.

3. Schumer M, Xu C, Powell DL, Durvasula A, Skov L, Holland C, et al. Natural selection interacts with recombination to shape the evolution of hybrid genomes. Science. 2018;360:656–60.

4. Abbott R, Albach D, Ansell S, Arntzen JW, Baird SJE, Bierne N, et al. Hybridization and speciation*. J Evol Biol. 2013;26:229–46.

5. Gompert Z, Parchman TL, Buerkle CA. Genomics of isolation in hybrids. Philosophical Transactions Royal Soc B Biological Sci. 2012;367:439–50.

6. Poikela N, Laetsch DR, Kankare M, Hoikkala A, Lohse K. Experimental introgression in Drosophila: Asymmetric postzygotic isolation associated with chromosomal inversions and an incompatibility locus on the X chromosome. Mol Ecol. 2023;32:854–66.

7. Cutter AD. The polymorphic prelude to Bateson–Dobzhansky–Muller incompatibilities. Trends Ecol Evol. 2012;27:209–18.

8. Langmore NE, Grealy A, Noh H-J, Medina I, Skeels A, Grant J, et al. Coevolution with hosts underpins speciation in brood-parasitic cuckoos. Science. 2024;384:1030–6.

9. Orr HA. Dobzhansky, Bateson, and the Genetics of Speciation. Genetics. 1996;144:1331–5.

10. Presgraves DC. Evaluating genomic signatures of “the large X-effect” during complex speciation. Mol Ecol. 2018;27:3822–30.

11. Kimura M. Evolutionary Rate at the Molecular Level. Nature. 1968;217:624–6.

12. Brevet M, Lartillot N. Reconstructing the history of variation in effective population size along phylogenies. Genome Biol Evol. 2021;13:evab150-.

13. Ohta T. eLS. 2013;

14. Cheng JY, Stern AJ, Racimo F, Nielsen R. Detecting Selection in Multiple Populations by Modeling Ancestral Admixture Components. Mol Biol Evol. 2021;39:msab294.

15. Runge J-N, Lindholm AK. Carrying a selfish genetic element predicts increased migration propensity in free-living wild house mice. Proc Royal Soc B. 2018;285:20181333.

16. Lynch M, Ackerman MS, Gout J-F, Long H, Sung W, Thomas WK, et al. Genetic drift, selection and the evolution of the mutation rate. Nat Rev Genet. 2016;17:704–14.

17. Zhang X-S, Hill WG. Genetic variability under mutation selection balance. Trends Ecol Evol. 2005;20:468–70.

18. Goyal S, Balick DJ, Jerison ER, Neher RA, Shraiman BI, Desai MM. Dynamic Mutation–Selection Balance as an Evolutionary Attractor. Genetics. 2012;191:1309–19.

19. Kuhlwilm M, Han S, Sousa VC, Excoffier L, Marques-Bonet T. Ancient admixture from an extinct ape lineage into bonobos. Nat Ecol Evol. 2019;3:957–65.

20. White MA, Stubbings M, Dumont BL, Payseur BA. Genetics and Evolution of Hybrid Male Sterility in House Mice. Genetics. 2012;191:917–34.

21. Schield DR, Scordato ESC, Smith CCR, Carter JK, Cherkaoui SI, Gombobaatar S, et al. Sex-linked genetic diversity and differentiation in a globally distributed avian species complex. Mol Ecol. 2021;30:2313–32.

22. Harris K, Nielsen R. The Genetic Cost of Neanderthal Introgression. Genetics. 2016;203:881–91.

23. Shchur V, Svedberg J, Medina P, Corbett-Detig R, Nielsen R. On the Distribution of Tract Lengths During Adaptive Introgression. G3 Genes Genomes Genetics. 2020;10:3663–73.

24. Sørensen EF, Harris RA, Zhang L, Raveendran M, Kuderna LFK, Walker JA, et al. Genome-wide coancestry reveals details of ancient and recent male-driven reticulation in baboons. Science. 2023;380:eabn8153.

25. Chiou KL, Bergey CM, Burrell AS, Disotell TR, Rogers J, Jolly CJ, et al. Genome-wide ancestry and introgression in a Zambian baboon hybrid zone. Mol Ecol. 2021;

26. Vilgalys TP, Fogel AS, Anderson JA, Mututua RS, Warutere JK, Siodi IL, et al. Selection against admixture and gene regulatory divergence in a long-term primate field study. Science. 2022;377:635–41.

27. Rogers J, Raveendran M, Harris RA, Mailund T, Leppälä K, Athanasiadis G, et al. The comparative genomics and complex population history of Papio baboons. Sci Adv. 2019;5:eaau6947.

28. Veller C, Edelman NB, Muralidhar P, Nowak MA. Recombination and selection against introgressed DNA. Evolution. 2023;77:1131–44.

29. Zeller E, Timmermann A. The evolving three-dimensional landscape of human adaptation. Sci Adv. 2024;10:eadq3613.

30. Vamosi SM, Schluter D. SEXUAL SELECTION AGAINST HYBRIDS BETWEEN SYMPATRIC STICKLEBACK SPECIES: EVIDENCE FROM A FIELD EXPERIMENT. Evolution. 1999;53:874–9.

31. Sánchez-Ramírez S, Weiss JG, Thomas CG, Cutter AD. Sex-specific and sex-chromosome regulatory evolution underlie widespread misregulation of inter-species hybrid transcriptomes. Biorxiv. 2020;2020.05.04.076505.

32. Larson EL, Vanderpool D, Sarver BAJ, Callahan C, Keeble S, Provencio LL, et al. The Evolution of Polymorphic Hybrid Incompatibilities in House Mice. Genetics. 2018;209:845–59.

33. Larson EL, Keeble S, Vanderpool D, Dean MD, Good JM. The composite regulatory basis of the large X-effect in mouse speciation. Mol Biol Evol. 2016;msw243.

34. Jolly CJ. Philopatry at the frontier: A demographically driven scenario for the evolution of multilevel societies in baboons (Papio). J Hum Evol. 2020;146:102819.

35. Zinner D, Wertheimer J, Liedigk R, Groeneveld LF, Roos C. Baboon phylogeny as inferred from complete mitochondrial genomes. Am J Phys Anthropol. 2013;150:133–40.

36. Skov L, Macià MC, Lucotte E, Cavassim MIA, Castellano D, Schierup MH, et al. Extraordinary selection on the human X chromosome associated with archaic admixture. Biorxiv. 2022;2022.09.19.508556.

37. Lucotte EA, Skov L, Jensen JM, Macià MC, Munch K, Schierup MH. Dynamic Copy Number Evolution of X- and Y-Linked Ampliconic Genes in Human Populations. Genetics. 2018;209:907–20.

38. Vicoso B, Charlesworth B. Effective Population Size and the Faster-X Effect: An Extended Model. Evolution. 2009;63:2413–26.

39. Meisel RP, Connallon T. The faster-X effect: integrating theory and data. Trends Genet. 2013;29:537–44.

40. Pool JE, Nielsen R. Inference of Historical Changes in Migration Rate From the Lengths of Migrant Tracts. Genetics. 2009;181:711–9.

41. Larson EL, Keeble S, Vanderpool D, Dean MD, Good JM. The composite regulatory basis of the large X-effect in mouse speciation. Molecular biology and evolution. 2016;msw243.

42. Rathje C, Johnson E, Drage D, Patinioti C, Silvestri G, Affara N, et al. Differential Sperm Motility Mediates the Sex Ratio Drive Shaping Mouse Sex Chromosome Evolution. Curr Biol. 2019;29:3692–3698.e4.

43. Hughes JF, Skaletsky H, Pyntikova T, Koutseva N, Raudsepp T, Brown LG, et al. Sequence analysis in Bos taurus reveals pervasiveness of X–Y arms races in mammalian lineages. Genome Res. 2020;

44. Skov L, Macià MC, Lucotte EA, Cavassim MIA, Castellano D, Schierup MH, et al. Extraordinary selection on the human X chromosome associated with archaic admixture. Cell Genom. 2023;100274.

45. Dutheil JY, Munch K, Nam K, Mailund T, Schierup MH. Strong Selective Sweeps on the X Chromosome in the Human-Chimpanzee Ancestor Explain Its Low Divergence. PLOS Genetics. 2015;11:e1005451.

46. Nam K, Munch K, Hobolth A, Dutheil J, Veeramah KR, Woerner AE, et al. Extreme selective sweeps independently targeted the X chromosomes of the great apes. Proceedings of the National Academy of Sciences. 2015;112:6413–8.

47. Nam K, Munch K, Mailund T, Nater A, Greminger M, Krützen M, et al. Evidence that the rate of strong selective sweeps increases with population size in the great apes. Proceedings of the National Academy of Sciences. 2017;114:201605660.

48. Kopp GH, Sithaldeen R, Trede F, Grathwol F, Roos C, Zinner D. A Comprehensive Overview of Baboon Phylogenetic History. Genes. 2023;14:614.

49. Vilgalys TP, Fogel AS, Mututua RS, Warutere JK, Siodi L, Anderson JA, et al. Selection against admixture and gene regulatory divergence in a long-term primate field study. Biorxiv. 2021;2021.08.19.456711.

50. Hellenthal G, Busby GBJ, Band G, Wilson JF, Capelli C, Falush D, et al. A Genetic Atlas of Human Admixture History. Science. 2014;343:747–51.

51. Wangkumhang P, Greenfield M, Hellenthal G. An efficient method to identify, date and describe admixture events using haplotype information. Biorxiv. 2021;2021.08.12.455263.

52. Kuderna LFK, Gao H, Janiak MC, Kuhlwilm M, Orkin JD, Bataillon T, et al. A global catalog of whole-genome diversity from 233 primate species. Science. 2023;380:906–13.

53. Liu KJ, Steinberg E, Yozzo A, Song Y, Kohn MH, Nakhleh L. Interspecific introgressive origin of genomic diversity in the house mouse. Proc Natl Acad Sci. 2015;112:196–201.

54. Maples BK, Gravel S, Kenny EE, Bustamante CD. RFMix: A Discriminative Modeling Approach for Rapid and Robust Local-Ancestry Inference. Am J Hum Genetics. 2013;93:278–88.

55. Massarat AR, Lamkin M, Reeve C, Williams AL, D’Antonio M, Gymrek M. Haptools: a toolkit for admixture and haplotype analysis. Bioinformatics. 2023;39:btad104.

56. Munch K, Nam K, Schierup MH, Mailund T. Selective Sweeps across Twenty Millions Years of Primate Evolution. Mol Biol Evol. 2016;33:3065–74.

57. Rivas-González I, Rousselle M, Li F, Zhou L, Dutheil JY, Munch K, et al. Pervasive incomplete lineage sorting illuminates speciation and selection in primates. Science. 2023;380:eabn4409.

58. Liang M, Nielsen R. The Lengths of Admixture Tracts. Genetics. 2014;197:953–67.

59. Dias-Alves T, Mairal J, Blum MGB. Loter: A Software Package to Infer Local Ancestry for a Wide Range of Species. Mol Biol Evol. 2018;35:2318–26.

60. Matheson J, Masel J. Background Selection From Unlinked Sites Causes Nonindependent Evolution of Deleterious Mutations. Genome Biol Evol. 2024;16:evae050.

61. Spence JP, Song YS. Inference and analysis of population-specific fine-scale recombination maps across 26 diverse human populations. Sci Adv. 2019;5:eaaw9206.

62. Kamm JA, Spence JP, Chan J, Song YS. Two-Locus Likelihoods Under Variable Population Size and Fine-Scale Recombination Rate Estimation. Genetics. 2016;203:1381–99.

63. Ronquist F, Kudlicka J, Senderov V, Borgström J, Lartillot N, Lundén D, et al. Universal probabilistic programming offers a powerful approach to statistical phylogenetics. Commun Biology. 2021;4:244.

64. Terhorst J, Kamm JA, Song YS. Robust and scalable inference of population history from hundreds of unphased whole genomes. Nat Genet. 2017;49:303–9.

65. Wall JD, Robinson JA, Cox LA. High-Resolution Estimates of Crossover and Noncrossover Recombination from a Captive Baboon Colony. Genome Biol Evol. 2022;14:evac040.

66. Pool JE, Nielsen R. POPULATION SIZE CHANGES RESHAPE GENOMIC PATTERNS OF DIVERSITY. Evolution. 2007;61:3001–6.

67. Charlesworth B, Jensen JD. Effects of Selection at Linked Sites on Patterns of Genetic Variability. Annu Rev Ecol, Evol, Syst. 2021;52:177–97.

68. Ellegren H, Galtier N. Determinants of genetic diversity. Nat Rev Genet. 2016;17:422–33.

69. Burri R, Nater A, Kawakami T, Mugal CF, Olason PI, Smeds L, et al. Linked selection and recombination rate variation drive the evolution of the genomic landscape of differentiation across the speciation continuum of Ficedula flycatchers. Genome Res. 2015;25:1656–65.

70. Nordborg M, Charlesworth B, Charlesworth D. The effect of recombination on background selection*. Genet Res. 1996;67:159–74.

71. Pool JE, Nielsen R. POPULATION SIZE CHANGES RESHAPE GENOMIC PATTERNS OF DIVERSITY. Evolution. 2007;61:3001–6.

72. Wu FL, Strand AI, Cox LA, Ober C, Wall JD, Moorjani P, et al. A comparison of humans and baboons suggests germline mutation rates do not track cell divisions. Plos Biol. 2020;18:e3000838.

73. Alberts SC, Watts HE, Altmann J. Queuing and queue-jumping: long-term patterns of reproductive skew in male savannah baboons, Papio cynocephalus. Anim Behav. 2003;65:821–40.

74. Alberts SC, Buchan JC, Altmann J. Sexual selection in wild baboons: from mating opportunities to paternity success. Anim Behav. 2006;72:1177–96.

75. Alberts SC. Social influences on survival and reproduction: Insights from a long-term study of wild baboons. J Anim Ecol. 2019;88:47–66.

76. Städele V, Roberts ER, Barrett BJ, Strum SC, Vigilant L, Silk JB. Male–female relationships in olive baboons (Papio anubis): Parenting or mating effort? J Hum Evol. 2019;127:81–92.

77. Kopp GH, Silva MJF da, Fischer J, Brito JC, Regnaut S, Roos C, et al. The Influence of Social Systems on Patterns of Mitochondrial DNA Variation in Baboons. Int J Primatol. 2014;35:210–25.

78. Hubisz MJ, Williams AL, Siepel A. Mapping gene flow between ancient hominins through demography-aware inference of the ancestral recombination graph. Plos Genet. 2020;16:e1008895.

79. McVicker G, Gordon D, Davis C, Green P. Widespread Genomic Signatures of Natural Selection in Hominid Evolution. Plos Genet. 2009;5:e1000471.

80. Chintalapati M, Moorjani P. Evolution of the mutation rate across primates. Curr Opin Genet Dev. 2020;62:58–64.

81. Conrad DF, Keebler JEM, DePristo MA, Lindsay SJ, Zhang Y, Casals F, et al. Variation in genome-wide mutation rates within and between human families. Nat Genet. 2011;43:712–4.

82. Oman M, Alam A, Ness RW. How Sequence Context-Dependent Mutability Drives Mutation Rate Variation in the Genome. Genome Biol Evol. 2022;14:evac032.

83. Hwang H-Y, Wang J. Effect of mutation mechanisms on variant composition and distribution in Caenorhabditis elegans. PLoS Comput Biol. 2017;13:e1005369.

84. Arbeithuber B, Betancourt AJ, Ebner T, Tiemann-Boege I. Crossovers are associated with mutation and biased gene conversion at recombination hotspots. Proc Natl Acad Sci. 2015;112:2109–14.

85. Hwang H-Y, Wang J. Effect of recombination on genetic diversity of Caenorhabditis elegans. Sci Rep. 2023;13:16425.

86. Fischer J, Higham JP, Alberts SC, Barrett L, Beehner JC, Bergman TJ, et al. Insights into the evolution of social systems and species from baboon studies. Elife. 2019;8:e50989.

87. Charlesworth B. The Effects of Deleterious Mutations on Evolution at Linked Sites. Genetics. 2012;190:5–22.

88. Speidel L, Forest M, Shi S, Myers SR. A method for genome-wide genealogy estimation for thousands of samples. Nat Genet. 2019;51:1321–9.

89. Breusch TS, Pagan AR. A Simple Test for Heteroscedasticity and Random Coefficient Variation. Econometrica. 1979;47:1287.

