## Supplementary for "Negative selection on baboon admixture is strongest on chromosome X"

### Simulation of admixture

To investigate the accuracy of RFMix[1] in detecting admixture, we performed a simulation using Haptools[2] with the simgenotype command. A pulse of 20 % Mikumi yellow baboons into Gog olive baboons was simulated, followed by 50 generations of recombination with no further admixture, and 10 individuals were extracted. Chromosome 8 was used as the reference VCF, and the recombination map inferred by Pyrho was used for the recombination landscape.

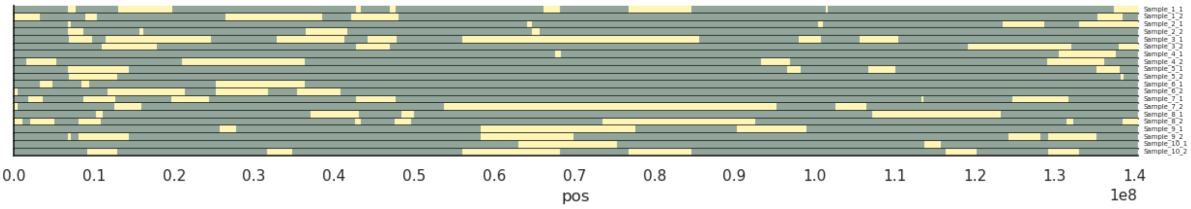


Simulation Figure 1: Visualisation of the simulated haplotypes from Haptools.

The resultant simulated haplotypes then depict a hypothetical admixture event between the almost pure Gog olive baboons and almost pure Mikumi yellow baboons and can be used to test whether RFMix is accurate and unbiased in its estimation of admixture. Simulation Figure 1 depicts the true ancestry of the various sections, as extracted from the breakpoints file generated by Haptools.

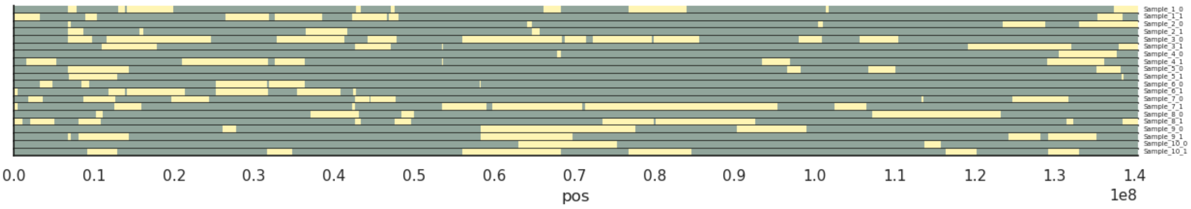


Simulation Figure 2: Visualisation of the inferred haplotypes from RFMix.

Using the same workflow as for the olive and yellow baboons to determine southern or northern ancestry, which is detailed in materials and methods, admixture was inferred with RFMix. To assess the accuracy of RFMix, admixture proportion in 100 kb windows was calculated. Visually, the RFMix inference is similar to simulated haplotypes (See Simulation Figure 2), but with one weakness: Very long fragments are broken up into smaller pieces by short stretches of opposite ancestry.

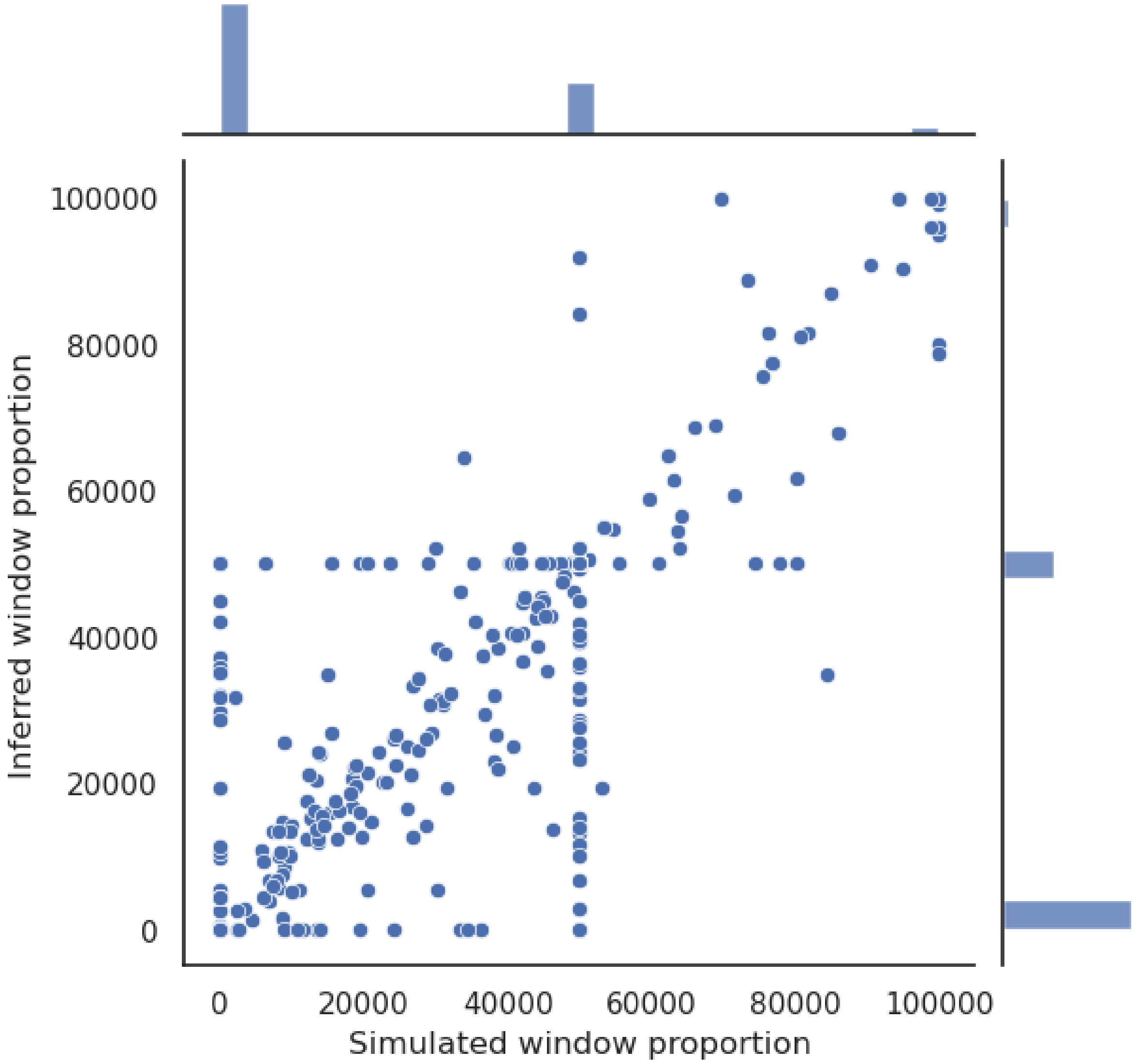


Simulation Figure 3: Simulated and inferred admixture in 100kb windows. Scatterplot of simulation and inference, with marginal histograms.

Pearson correlation for the simulation and inference is 98.9 %, and SpearmanR correlation for the simulation and inference is 96.4 %, with a p-value of 0 for both. See simulation Figure 3 for the distribution of values used for the correlations. There is no significant difference in the degree of inferred admixture under a Student's T-test (p-value 0.655). 17.4% MPA from southern baboons is inferred, and the true degree of MPA is 17,6%.

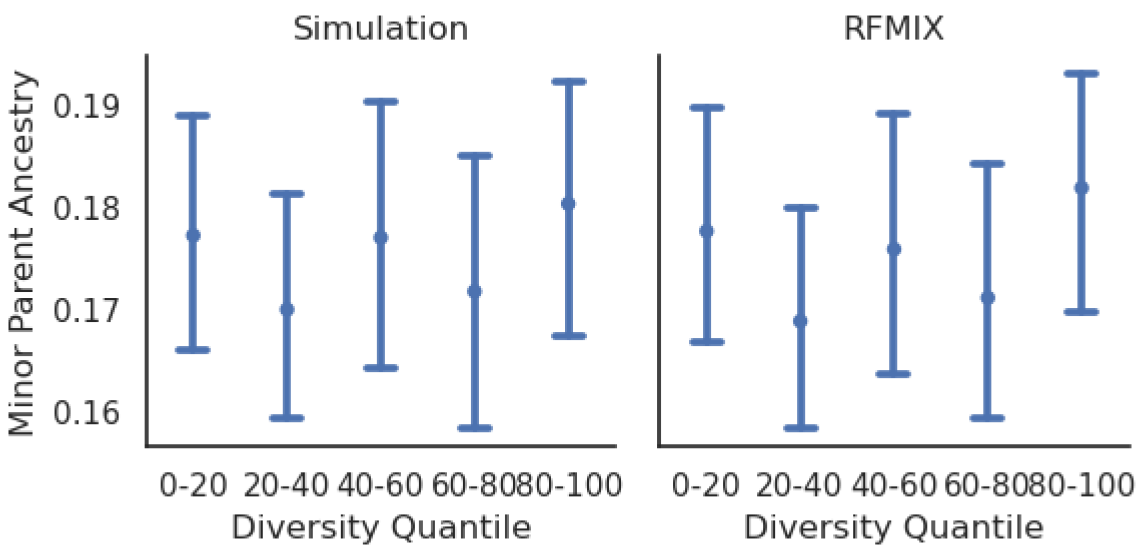


Simulation Figure 4: Minor Parent Ancestry for Simulation and RFMIX inference.

All Diversity Quantiles overlap the true admixture proportion, and the confidence intervals for each quantile are similar (Simulation Figure 4).

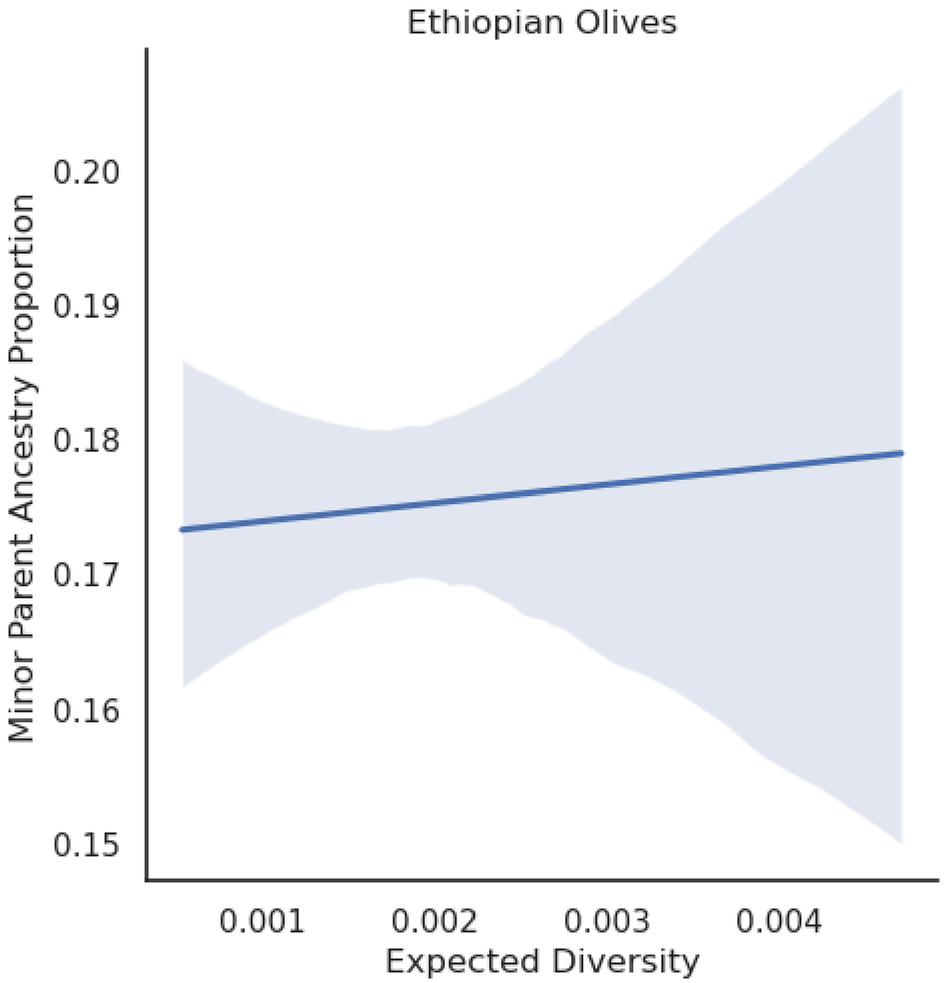


Simulation Figure 5: Minor Parent Ancestry regression.

A weighted linear regression, performed in the same way as the regressions for the autosomes and chromosome X in the main also shows no significant correlation between background diversity and inferred Minor Parent Ancestry proportion (p-value 0.763), see Simulation Figure 5.

#### Distribution and regressions with outliers

Outliers can bias regressions, either due to errors in outlier regions, or due to outlier regions being significantly different from other regions due to actual biological dynamics which makes it differ from the rest of the genome.

Errors could for example be due to badly assembled regions which will have elevated diversity due to overcollapse, which is when a region which is thought to be only a single region is repeated one or more times across the genome, leading to an elevated density of SNPs. A callability mask will catch the most egregious regions of this kind, but if the region only is duplicated once, it will be difficult to impossible to discern only based on read depth.

On the other hand, high-recombination or high-diversity regions might also arise due to unique selective pressures, such as balancing selection or recombination hotspots. Recombination hotspots are relevant, but it is difficult to estimate them correctly, so the highest hotspots might be overestimated [3]. In addition, the process of linked selection does require recombination events, but the highest recombination regions do not necessarily confer a much greater benefit than intermediate-level recombination in removing linkage.

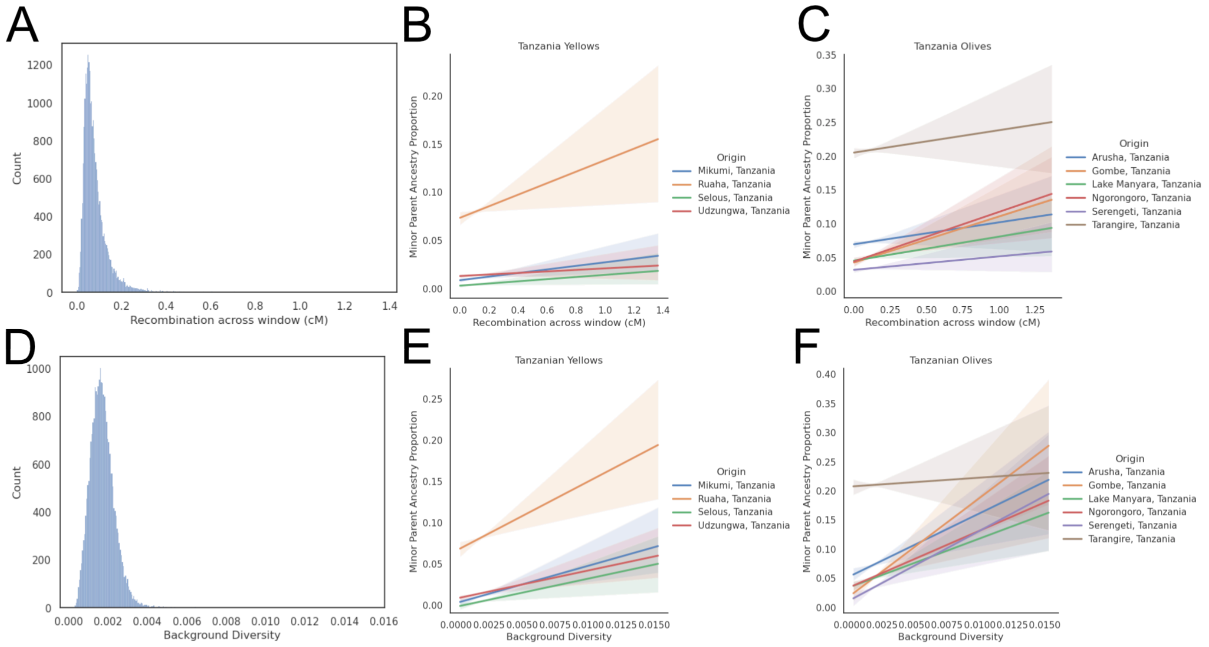


**Outlier Figure 1:** ***(A)*** *Distribution of recombination as measured in cM across the autosomes.* ***B-C)*** *Weighted Linear Regressions of the Tanzanian baboon populations with recombination.* ***(D)*** *Distribution of background diversity across the autosomes.* ***E-F)*** *Weighted Linear Regressions of the Tanzanian baboon populations with background diversity.*

Not removing outliers shows a large skew in both recombination and diversity (Outlier Figure 1A, 1D). Selous, Mikumi and Udzungwa have less steep slopes, while Ruaha has a steeper slope with recombination (Outlier Figure 1B, Outlier Table 1) compared to the case without removing outliers. Similarly, the results are also slightly different with Tanzanian olive baboons (Outlier Figure 1C), with some populations (Ngorongoro, Gombe and Lake Manyara) with a steeper slope, and others (Serengeti) with a less steep slope. Note that

Tarangire and Arusha baboons have a significant slope in this regression, unlike when outliers are removed. Lastly, Gog depicts the olive-hamadryas admixture case of Ethiopian olive baboons.

When leveraging background diversity (Outlier Figure 1D-1F, Outlier Table 2), the results are also heterogenous, but still with significant p-values for all cases presented as significant in the main article. Olive baboons in Arusha also have a significant slope with background diversity, but Tarangire baboons do not.

**Outlier Table 1:** Results for the weighted linear regression based on MPA and recombination rate.

| Origin | Intercept | Slope | Intercept P-value | Slope P-value | Intercept stderr | Slope stderr |
|---|---|---|---|---|---|---|
| Selous, Tanzania | 0.0032 | 0.0112 | 2.39e-19 | 1.82e-08 | 0.000355 | 0.00199 |
| Mikumi, Tanzania | 0.00885 | 0.0185 | 1.19e-213 | 1.59e-31 | 0.000284 | 0.00159 |
| Udzungwa, Tanzania | 0.0132 | 0.00792 | 6.55e-231 | 0.000554 | 0.000407 | 0.00229 |
| Ruaha, Tanzania | 0.0736 | 0.0598 | 0 | 5.46e-30 | 0.000933 | 0.00526 |
| Tarangire, Tanzania | 0.205 | 0.033 | 0 | 0.000329 | 0.00163 | 0.0092 |
| Arusha, Tanzania | 0.0694 | 0.0323 | 0 | 3.51e-07 | 0.00113 | 0.00634 |
| Ngorongoro, Tanzania | 0.0438 | 0.0733 | 0 | 4.33e-44 | 0.000934 | 0.00527 |
| Gombe, Tanzania | 0.0423 | 0.0681 | 0 | 4.14e-51 | 0.000803 | 0.00453 |
| Lake Manyara, Tanzania | 0.0452 | 0.0354 | 0 | 8.6e-18 | 0.000732 | 0.00412 |
| Serengeti, Tanzania | 0.0315 | 0.0199 | 0 | 1.18e-08 | 0.00062 | 0.00349 |
| Gog Woreda, Ethiopia | 0.0746 | 0.176 | 0 | 3.24e-93 | 0.00152 | 0.00857 |

**Outlier Table 2:** Results for the weighted linear regression based on MPA and background diversity.

| Origin | Intercept | Slope | Intercept P-value | Slope P-value | Intercept stderr | Slope stderr |
|---|---|---|---|---|---|---|
| Selous, Tanzania | 0.00209 | 1.13 | 0.000581 | 0.000591 | 0.000607 | 0.329 |
| Mikumi, Tanzania | 0.00599 | 2.54 | 1.5e-35 | 1.67e-22 | 0.000481 | 0.26 |
| Udzungwa, Tanzania | 0.00782 | 3.27 | 1.67e-25 | 7.92e-16 | 0.00075 | 0.406 |
| Ruaha, Tanzania | 0.0681 | 7.08 | 0 | 4.01e-14 | 0.00173 | 0.936 |
| Tarangire, Tanzania | 0.218 | -4.1 | 0 | 0.0132 | 0.00305 | 1.65 |
| Arusha, Tanzania | 0.0657 | 4.31 | 1.22e-211 | 0.000169 | 0.00212 | 1.15 |
| Ngorongoro, Tanzania | 0.0408 | 6.34 | 2.03e-125 | 8.39e-12 | 0.00171 | 0.928 |
| Gombe, Tanzania | 0.0298 | 11.3 | 1.89e-94 | 5.58e-47 | 0.00145 | 0.784 |
| Lake Manyara, Tanzania | 0.0409 | 4.99 | 9.55e-199 | 1.28e-11 | 0.00136 | 0.737 |
| Serengeti, Tanzania | 0.0225 | 6.41 | 1.32e-93 | 4.55e-27 | 0.0011 | 0.595 |
| Gog Woreda, Ethiopia | 0.0322 | 31.9 | 4.38e-35 | 9.06e-114 | 0.0026 | 1.41 |

### Positive selection scan

Positive selection decreases nearby diversity in the genome due to linked selection. In addition, Linkage Disequilibrium in the area increases due to the swept haplotype being a relatively recent common ancestor. Relate[78] can detect this signal by building trees for genome sections and detecting if rapid coalescences occur after a mutation is present. The patterns of adjacent mutations with no observed recombination events allow for estimating the time to the most recent common ancestor of the derived lineages. Sample sizes in individual populations are not large enough to allow for selective sweep inference using Relate, and all olives in Tanzania are therefore grouped. All yellows in Tanzania have a sample size that is too low and show no log p-values above 7.5.

There is no evidence of adaptive introgressions in Tanzanian olive baboons. Of the 150 100 kb windows with evidence for positive selection inferred by Relate (log-p-value >= 7.5) on the autosomes (See Supplementary Figure 14A-B), there is, on average, 32.5 % less admixture (corresponding to 1.5 percentage points less admixture in the swept regions, from 6.6 % to 5.1 % admixture). Using Welch's T-test, there is a significant difference in the autosomes between admixture percentage in swept regions and those with no detected sweeps (p-value 1.68e-13), indicating that the recent selective sweeps in olive baboons primarily happen in regions depleted of MPA from southern baboons and that adaptive introgressions either are too old to detect for Tanzanian olives or not present. Using a T-test for chromosome X (p-value threshold lowered to log >= 6), there is no significant difference (p-value 0.551) between levels of admixture in swept regions and regions without sweeps (See Supplementary Figure 14C).

### Commands used for analysis

Bcftools filtering:
bcftools filter -e "(GT='./.') | (GT='het' & FMT/AD[*:*] < 3 ) | FMT/DP <= $min_cov | FMT/DP >= $max_cov | FMT/GQ <= 30

SMC++ run:
--timepoints 10 10e6 --spline piecewise \ --ftol 1e-3 --em-iterations 10.

Pyrho maketable:
--decimate_rel_tol 0.25 –approx

Pyrho optimize:
--blockpenalty 10,25,50,100 and –windowsize 10,25,50,100.

RFMix:
-e 3 -G 100 --reanalyze-reference

Statsmodels regressions:
MPA ~ recombination_rate
MPA ~ background_diversity
MPA ~ recombination_rate + background_diversity
MPA ~ normalized_diversity * chrom_type

Supplementary Figures

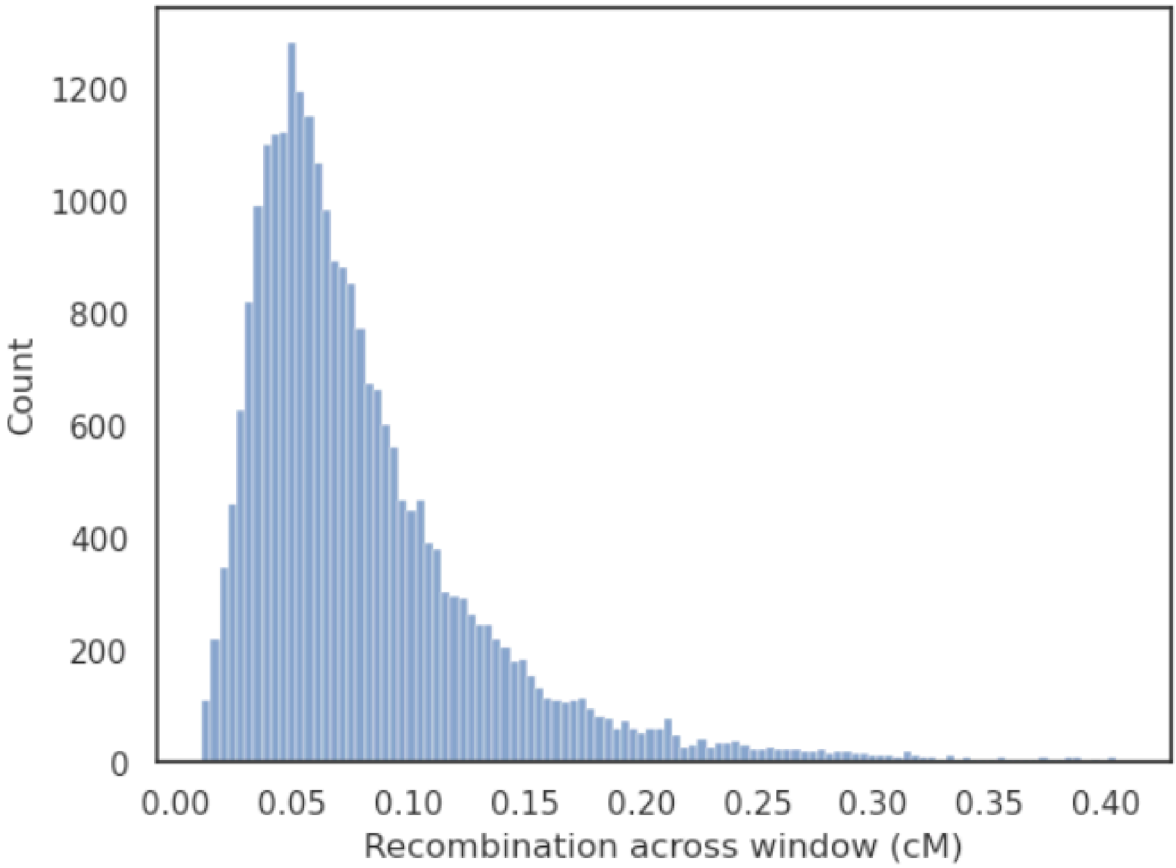


**Supplementary Figure 1:** Distribution of recombination as measured in cM across the autosomes after filtering the 0.5 % high and low outliers.

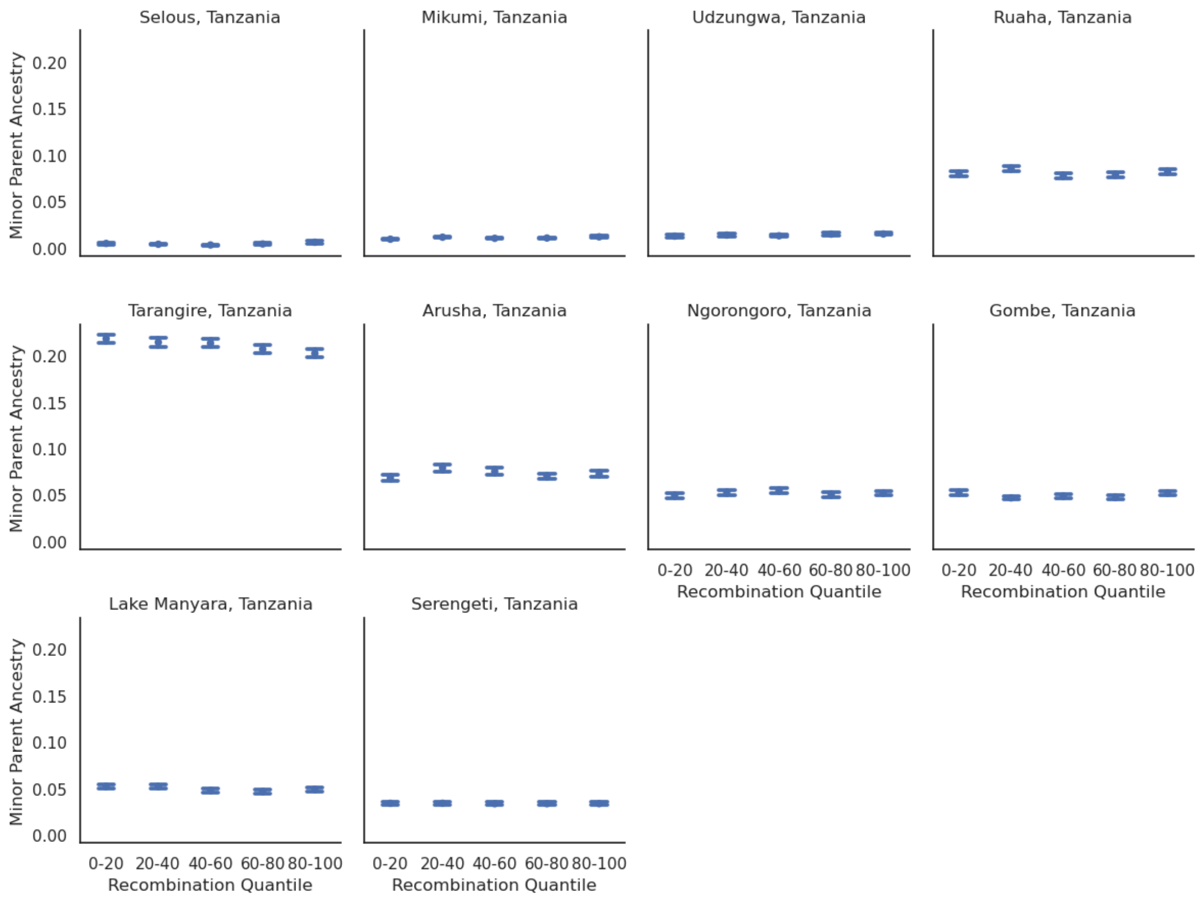


**Supplementary Figure 2:** Quintile distribution of all sampled Tanzanian populations based on recombination rate and Minor Parent Ancestry on the autosome.

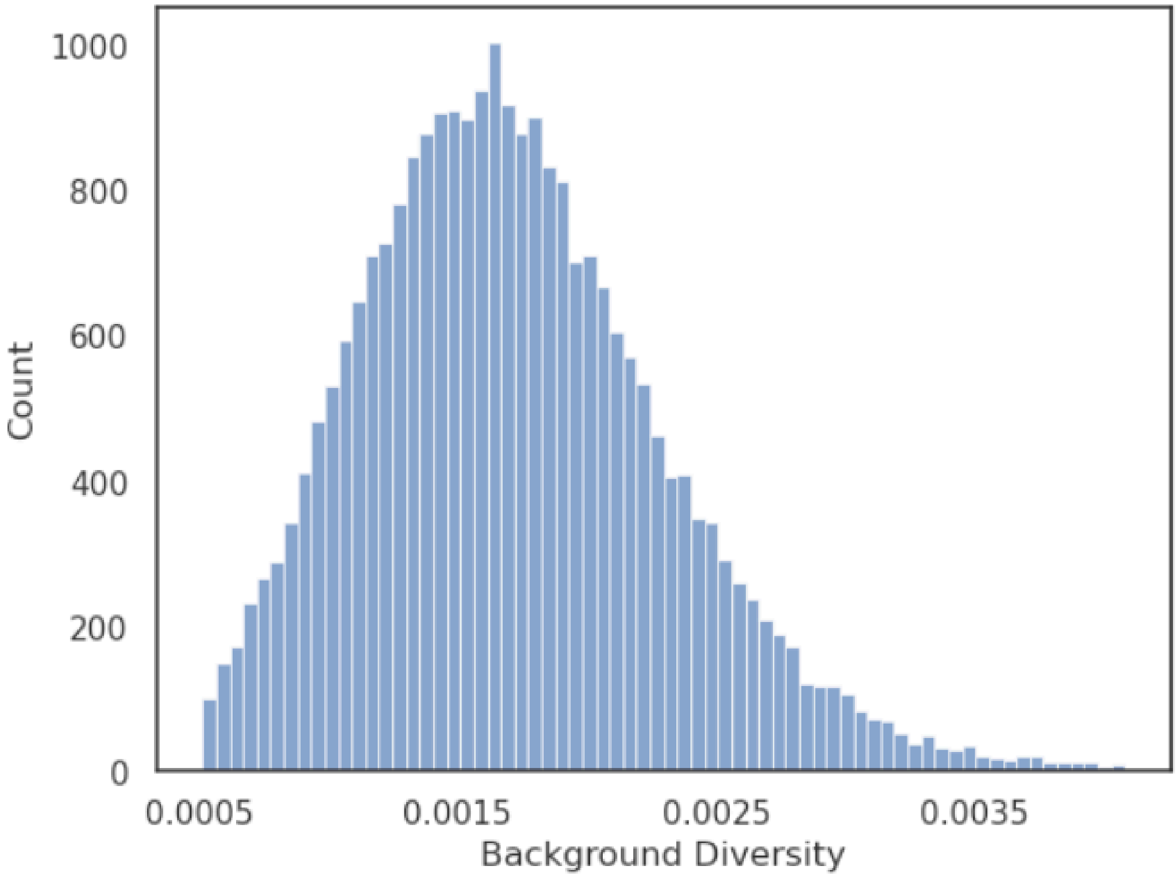


**Supplementary Figure 3:** Distribution of Background Diversity across the autosomes after filtering the 0.5 % high and low outliers. Background diversity on the autosomes has a mean of 0.00171 and a median of 0.00165

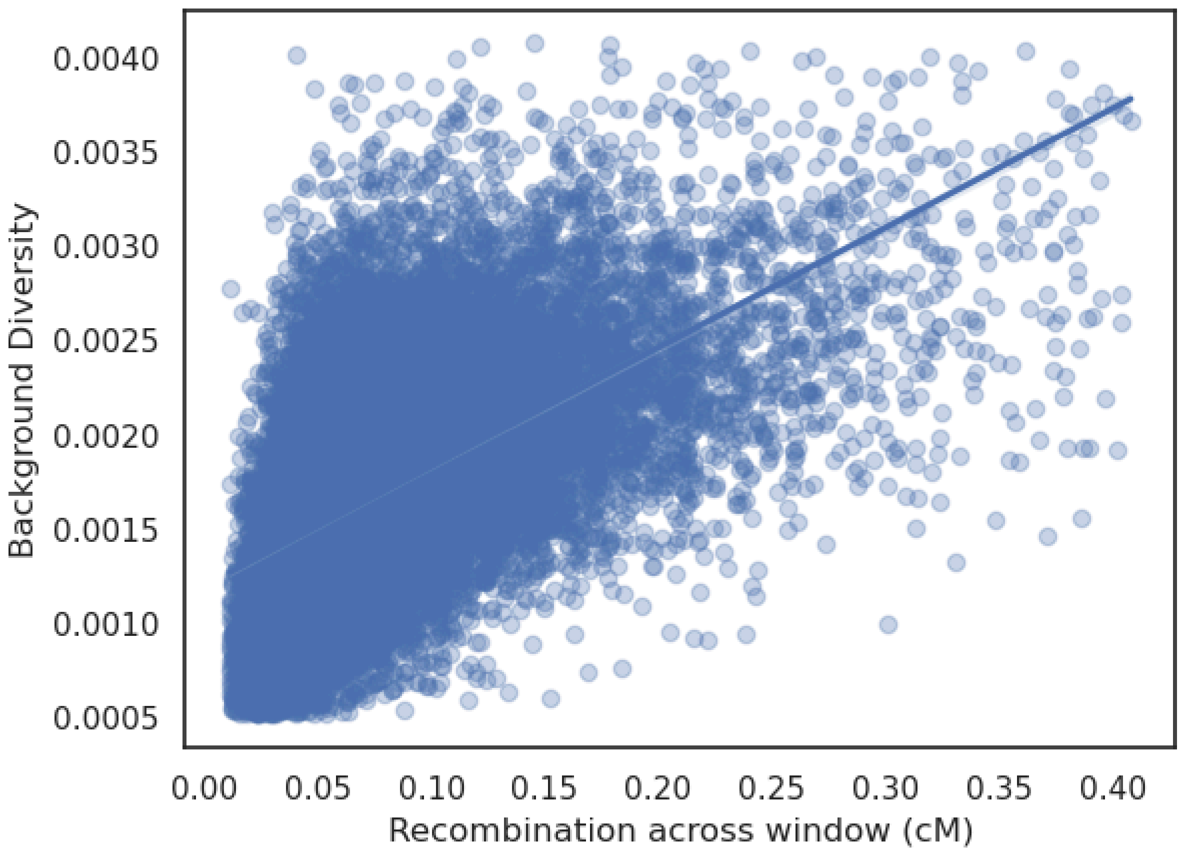


**Supplementary Figure 4:** Correlation between recombination and Background Diversity after filtering outliers 0.5 % high and low recombination rate and background diversity windows on the autosome.

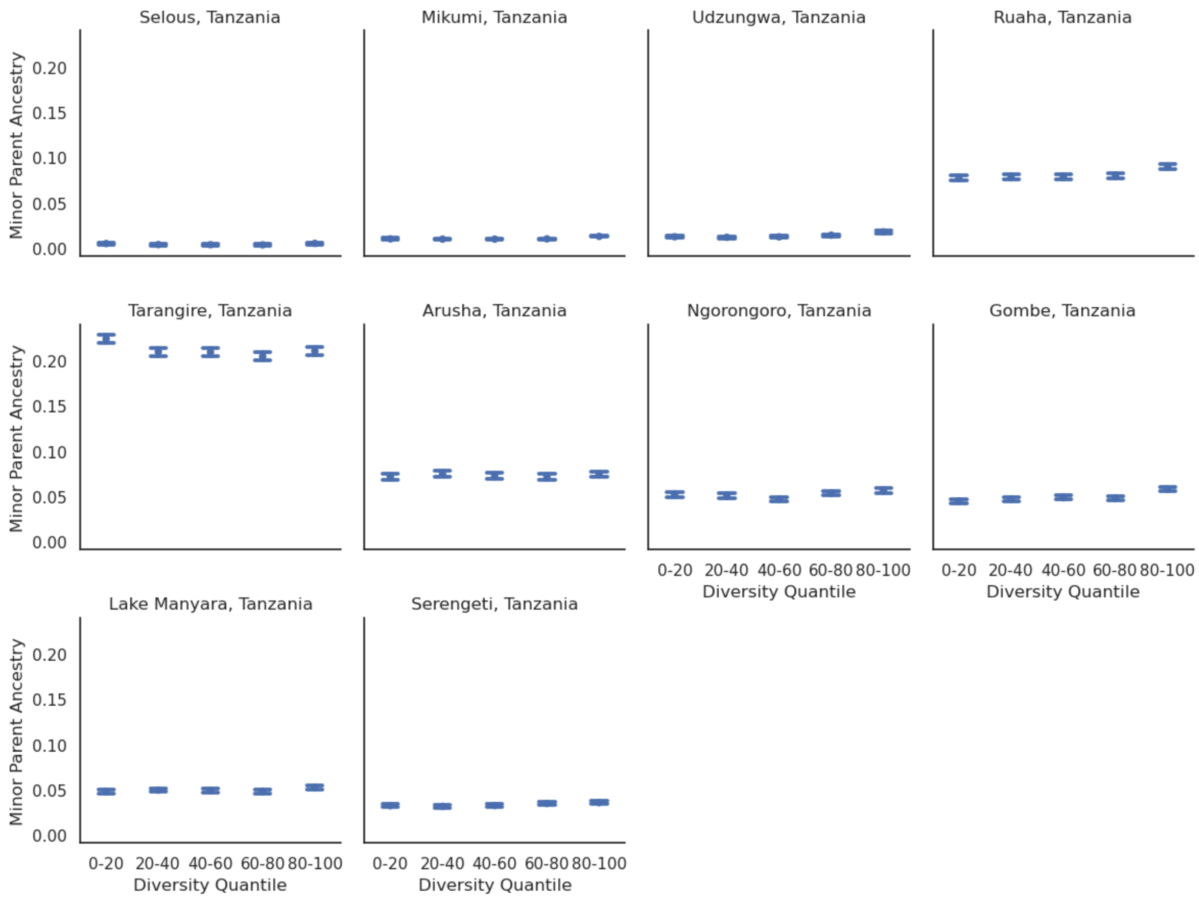


**Supplementary Figure 5:** Quintile distribution of all sampled Tanzanian populations based on Background Diversity and Minor Parent Ancestry on the autosomes.

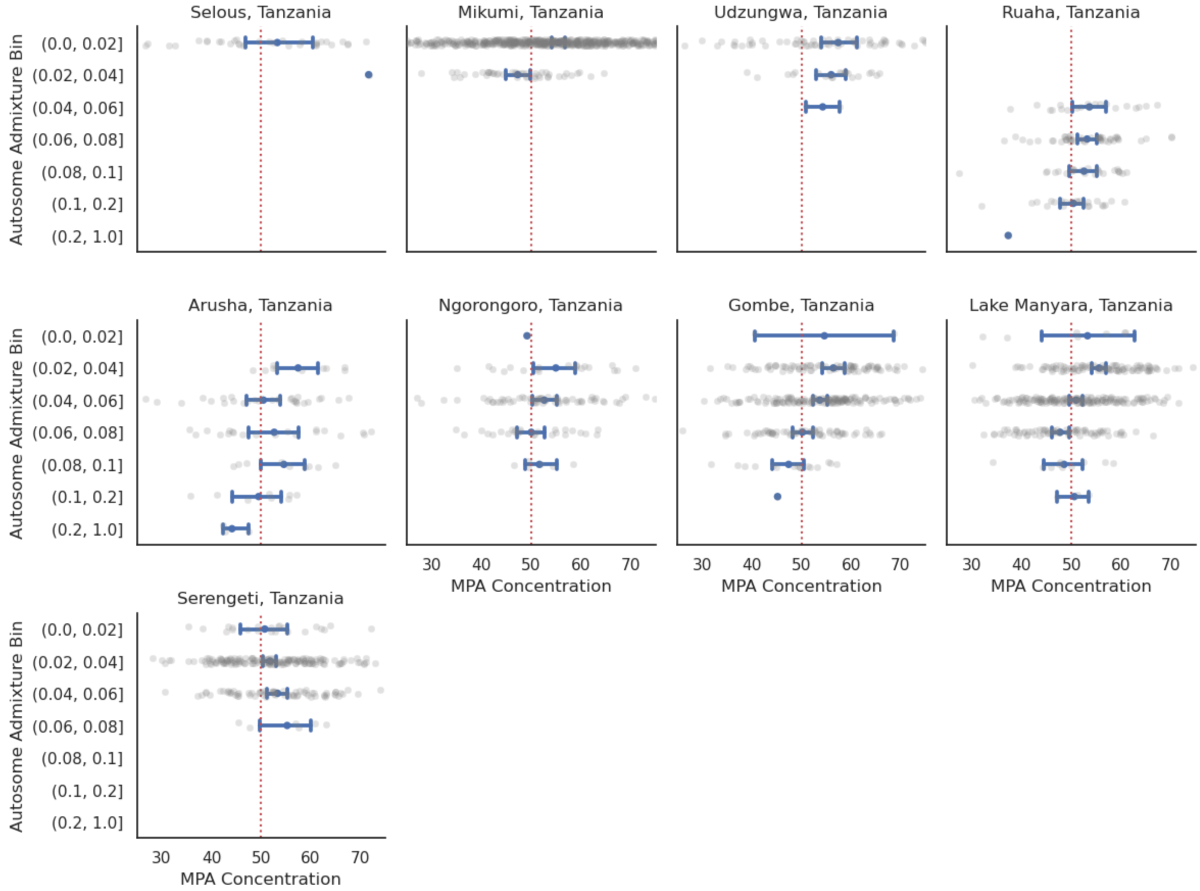


**Supplementary Figure 6:** Minor Parent Ancestry concentration stratified based on admixture level. MPA concentration is defined as which percentage of admixture is found in the most half of the chromosome with the highest background diversity. The red line represents the expectation of 50 % admixture in the 50 % of the chromosome with the highest diversity. Grey dots represent individual chromosomes, and the blue error bar represents the 95 % confidence interval. Note that the first five bins are equally sized until 10 % admixture, and the last two bins correspond to admixture levels above 10 % and 20 % admixture, respectively.

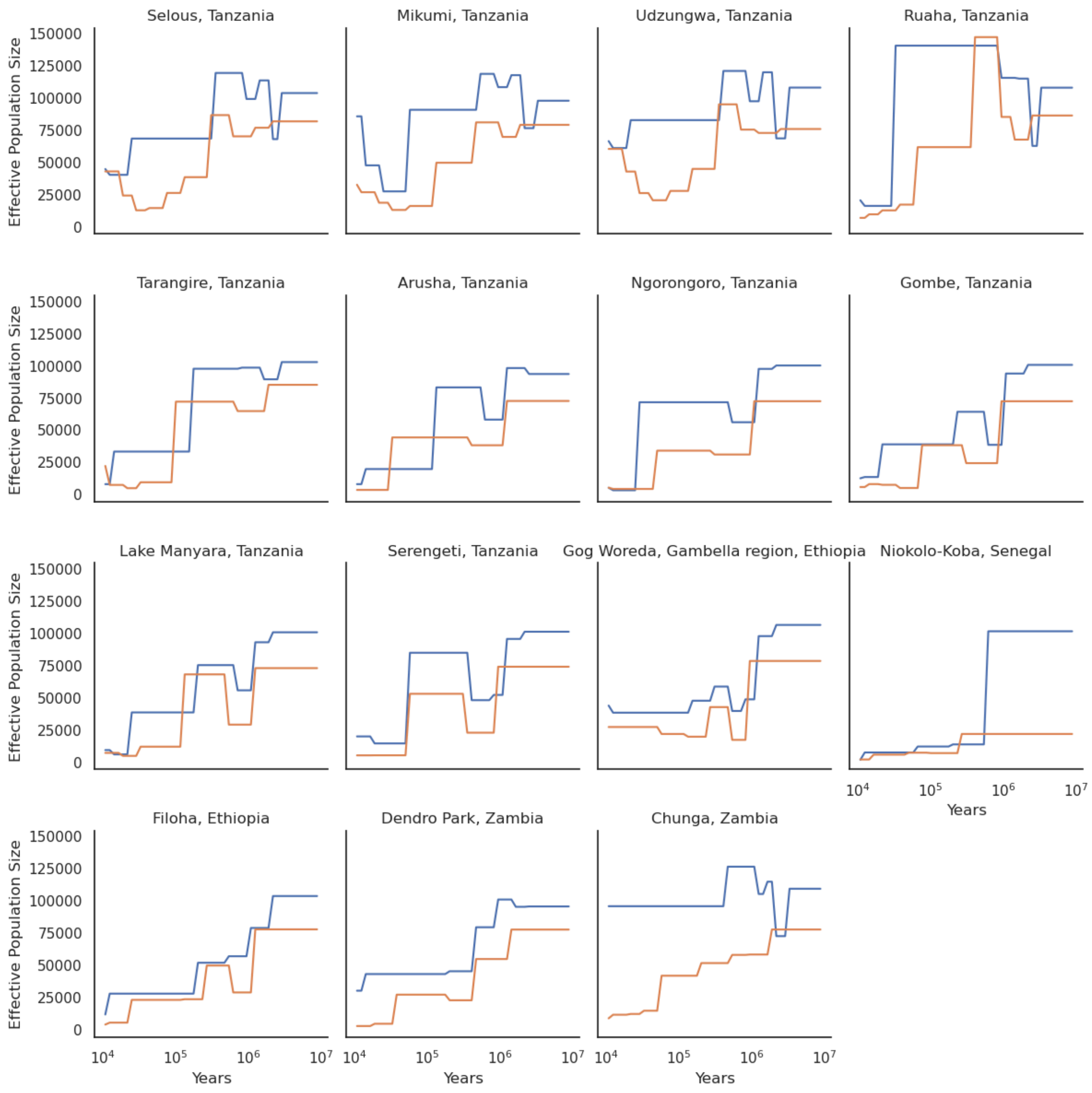


**Supplementary Figure 7:** Population size history for all 15 populations used in this study as inferred by SMC++. Selous to Ruaha are yellow baboon populations, Tarangire to Gog are olive baboons, Niokolo-Koba is guinea baboons, Filoha is hamadryas baboons, Dendro Park is chacma baboons, and Chunga is kinda baboons. All populations except Niokolo-Koba recover to an ancestral state with approximately 100000 Ne for autosomes and 75000 for chromosome X.

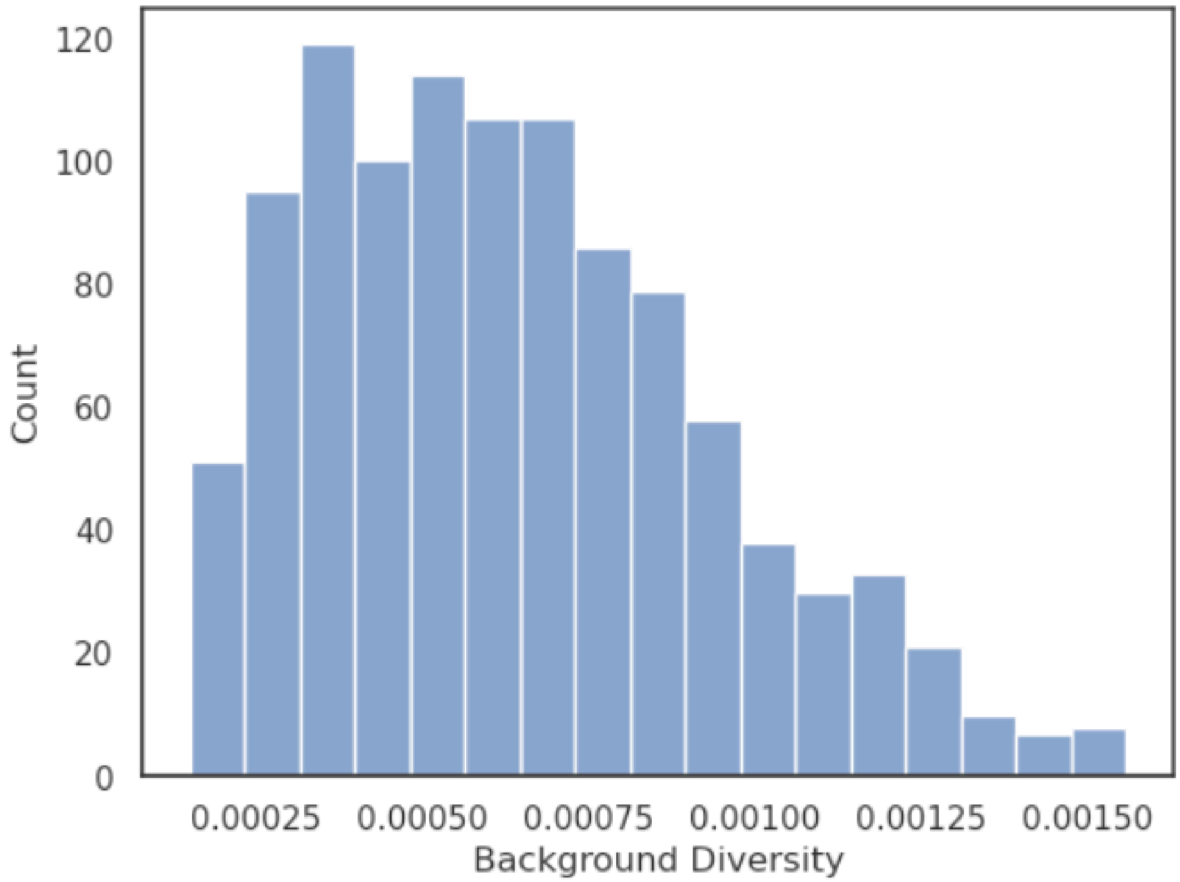


**Supplementary Figure 8:** Distribution of background diversity across chromosome X after filtering the 0.5 % high and low outliers.

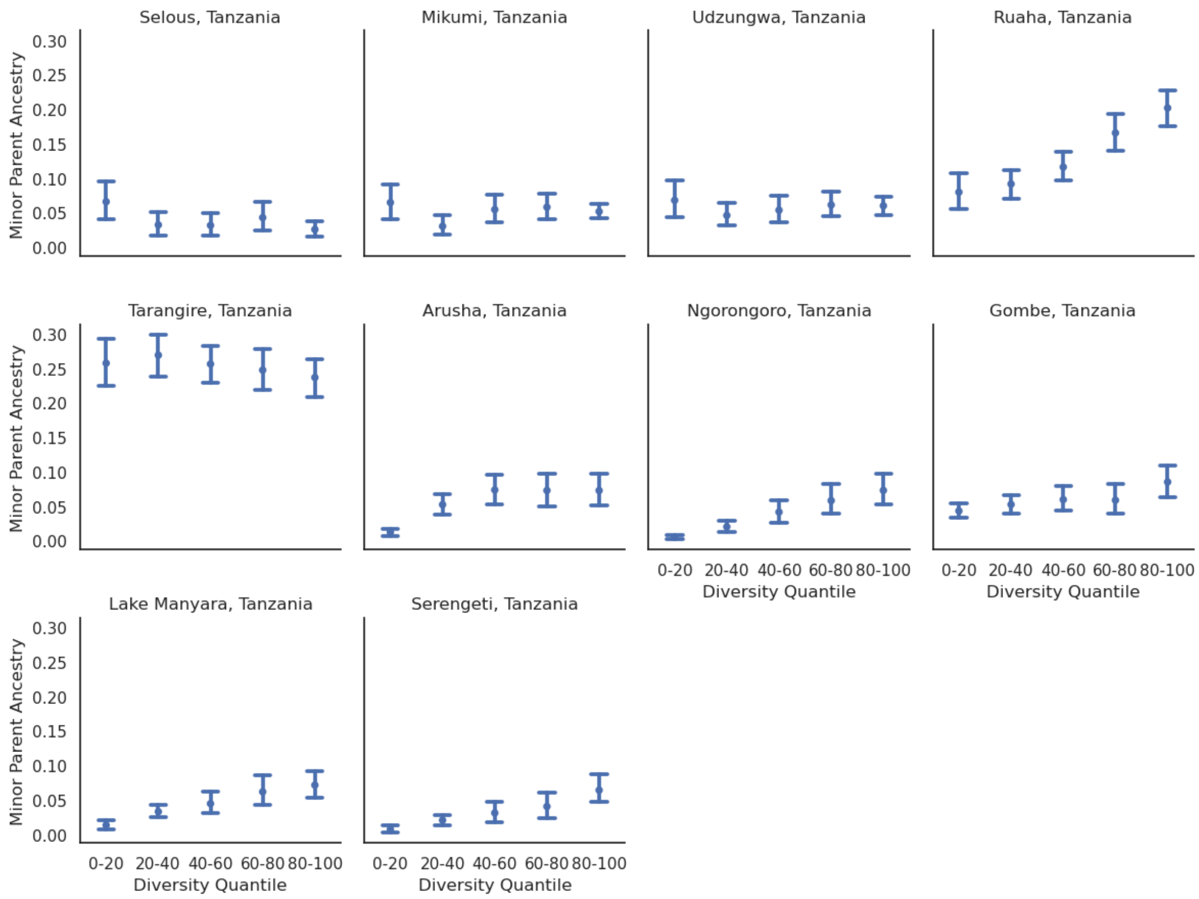


**Supplementary Figure 9:** Quintile distribution of all sampled Tanzanian populations based on Background Diversity and Minor Parent Ancestry for chromosome X.

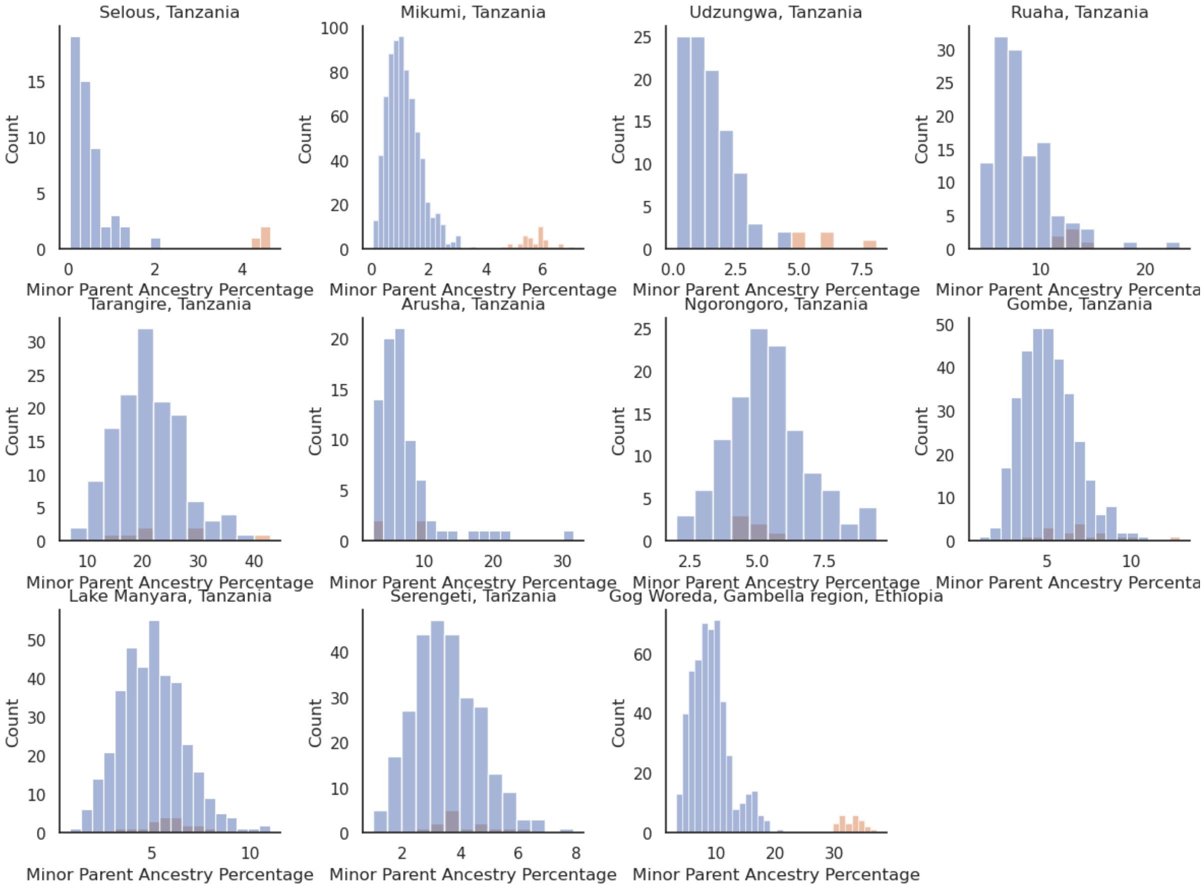


**Supplementary Figure 10:** Depicts the admixture percentage of each chromosome, with the autosomes (1 to 20) depicted in blue and chromosome X depicted in orange. Each individual therefore contributes 20 autosomal counts and 1 chromosome X count. Gog Woreda depicts the Hamadryas admixture case

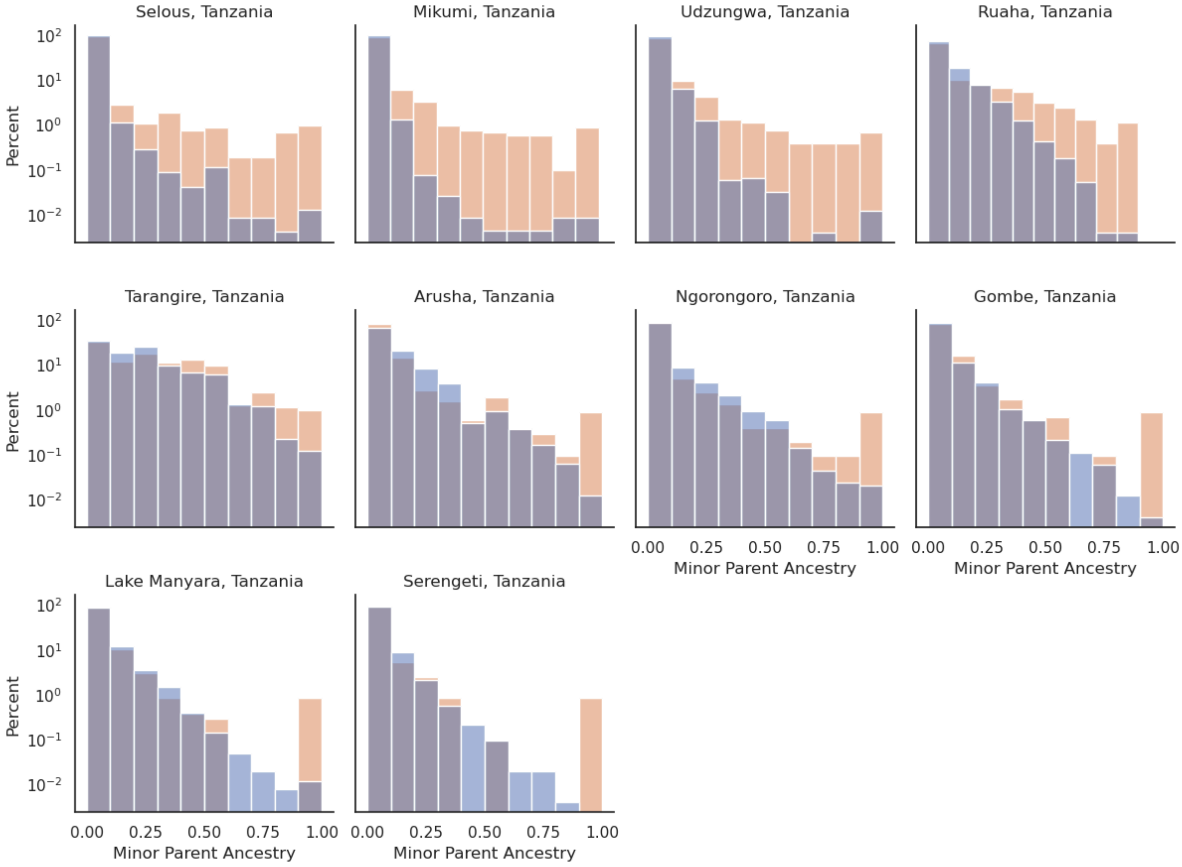


**Supplementary Figure 11:** Minor Parent Ancestry percentage distribution. Blue denotes autosomal frequency and orange denotes chromosome X. Note that it is log scaled, as all populations have a majority of their windows without any MPA.

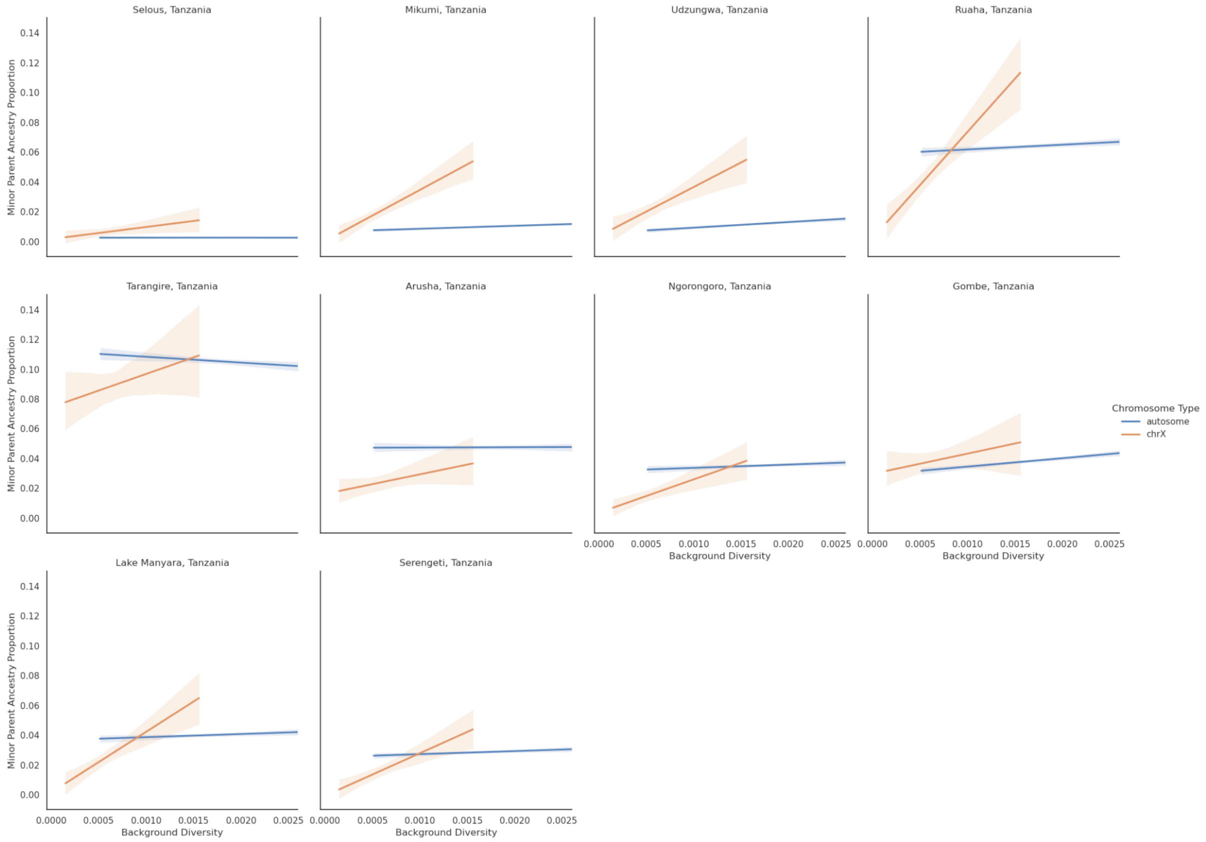


**Supplementary Figure 12:** Truncated plot of Expected Diversity and MPA on the autosomes and chromosome X for Tanzanian olive and yellow baboons, after filtering out all windows with more than 25 % MPA.

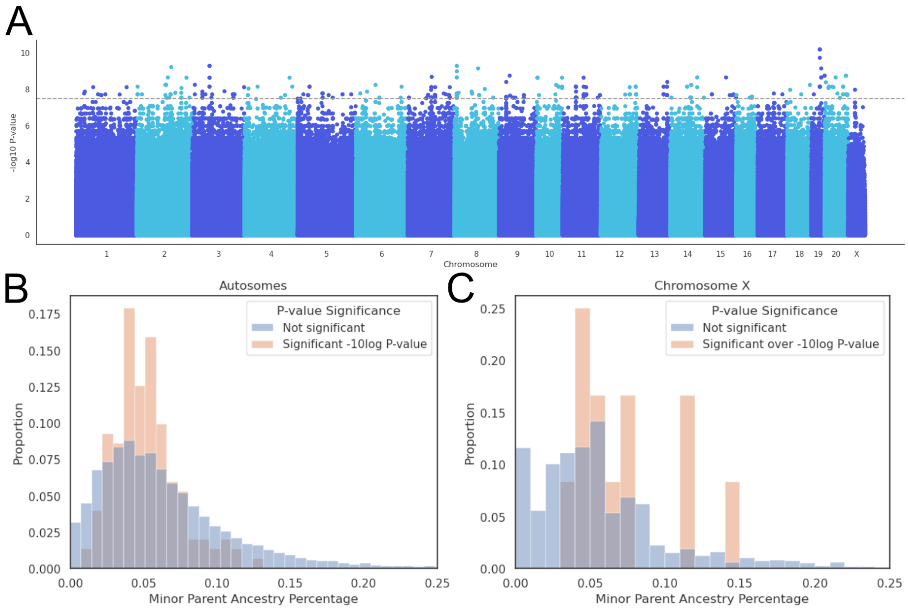


**Supplementary Figure 13**: **A)** Manhattan plot of -log p-values. **B)** Histogram of MPA proportion of significant p-values and MPA proportion of not significant p-values on the autosomes. **C)** Histogram of MPA proportion of significant p-values and MPA proportion of not significant p-values on chromosome X.

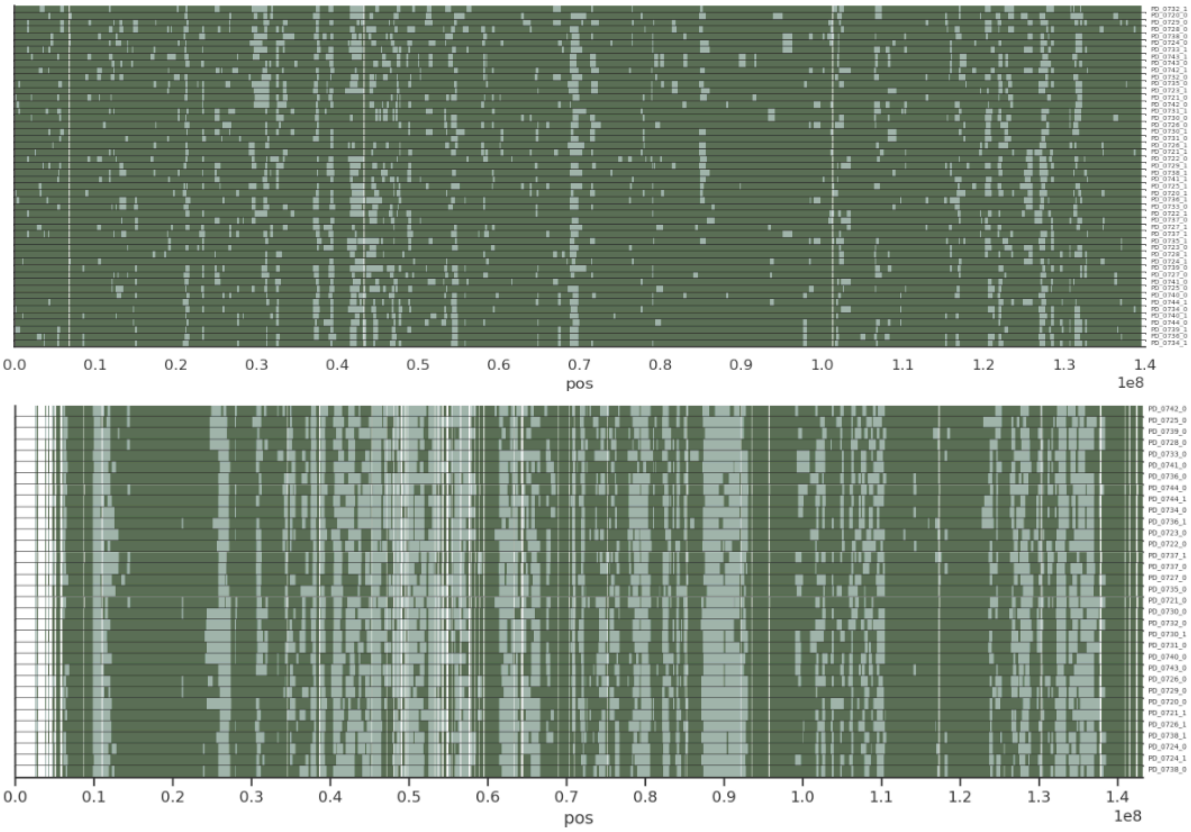


**Supplementary Figure 14:** RFMix paintings of chrX and chr8 for the Gog olives. The haplotypes are ordered based on their similarity using UPGMA, and cluster based on the sampling location. Windows with less than 75 % callability are filtered out. Note that there are fewer haplotypes for Chromosome X, as males only contain 1 copy instead of 2.

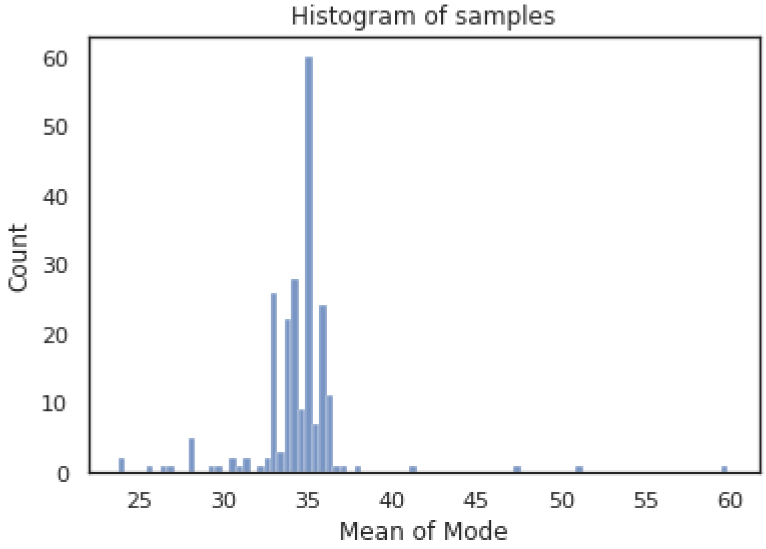


**Supplementary Figure 15:** Every chromosomes mode callability depth was calculated, and then averaged per individual. Most individuals are distributed around 35X depth.

#### Supplementary Tables

**Supplementary Table 1:** Overview of autosomal and X-linked admixture per population, as well as sample number of males and females.

| Origin | Autosomal admixture % | Chromosome X admixture % |
|---|---|---|
| Selous, Tanzania | 0.361 | 4.52 |
| Mikumi, Tanzania | 1.04 | 5.72 |
| Udzungwa, Tanzania | 1.35 | 6.14 |
| Ruaha, Tanzania | 8.04 | 12.9 |
| Tarangire, Tanzania | 20.9 | 25 |
| Arusha, Tanzania | 7.31 | 6.2 |
| Ngorongoro, Tanzania | 5.18 | 4.2 |
| Gombe, Tanzania | 4.98 | 6.31 |
| Lake Manyara, Tanzania | 4.98 | 5.05 |
| Serengeti, Tanzania | 3.4 | 3.48 |
| Gog, Ethiopia | 0.0654 | 1.3 |

**Supplementary Table 2:** Breusch-Pagan Lagrange Multiplier test for heteroscedasticity for the distribution of recombination and Minor Parent Ancestry on the autosomes.

| Lagrange Multiplier | Lagrange Multiplier P-value | F-statistic | F-statistic P-value | Origin |
|---|---|---|---|---|
| 0.000847 | 0.977 | 0.000847 | 0.977 | Selous, Tanzania |
| 6.37 | 0.0116 | 6.37 | 0.0116 | Mikumi, Tanzania |
| 0.0245 | 0.876 | 0.0245 | 0.876 | Udzungwa, Tanzania |
| 12.5 | 0.000414 | 12.5 | 0.000414 | Ruaha, Tanzania |
| 0.0695 | 0.792 | 0.0695 | 0.792 | Tarangire, Tanzania |
| 0.0011 | 0.974 | 0.0011 | 0.974 | Arusha, Tanzania |
| 1.38 | 0.24 | 1.38 | 0.24 | Ngorongoro, Tanzania |
| 17.9 | 2.33e-05 | 17.9 | 2.32e-05 | Gombe, Tanzania |
| 4.06 | 0.0439 | 4.06 | 0.0439 | Lake Manyara, Tanzania |
| 10.1 | 0.00147 | 10.1 | 0.00147 | Serengeti, Tanzania |

**Supplementary Table 3:** Results for the weighted linear regressions of MPA against recombination rate.

| Origin | Intercept | Slope | Intercept P-value | Slope P-value | Intercept stderr | Slope stderr |
|---|---|---|---|---|---|---|
| Selous, Tanzania | 0.00137 | 0.0281 | 0.00229 | 7.3e-18 | 0.00045 | 0.00327 |
| Mikumi, Tanzania | 0.00728 | 0.0325 | 1.27e-94 | 1.07e-37 | 0.000353 | 0.00253 |
| Udzungwa, Tanzania | 0.0107 | 0.0309 | 1.28e-97 | 6.38e-17 | 0.000509 | 0.00369 |

| | | | | | | |
|---|---|---|---|---|---|---|
| Ruaha, Tanzania | 0.0759 | 0.0414 | 0 | 8.29e-07 | 0.00116 | 0.00841 |
| Tarangire, Tanzania | 0.212 | -0.0327 | 0 | 0.027 | 0.00204 | 0.0148 |
| Arusha, Tanzania | 0.0717 | 0.00994 | 0 | 0.327 | 0.0014 | 0.0102 |
| Ngorongoro, Tanzania | 0.0491 | 0.0261 | 0 | 0.00177 | 0.00115 | 0.00836 |
| Gombe, Tanzania | 0.0428 | 0.0633 | 0 | 5.6e-19 | 0.000981 | 0.00711 |
| Lake Manyara, Tanzania | 0.0474 | 0.0142 | 0 | 0.0306 | 0.000903 | 0.00655 |
| Serengeti, Tanzania | 0.0306 | 0.0279 | 0 | 4.07e-07 | 0.00076 | 0.00551 |

**Supplementary Table 4:** Minor Parent Percentage in the lowest and highest 20% recombination quantile, as well as the relative and absolute difference on the autosomes.

| Origin | 0-20 Minor Parent Percentage | 80-100 Minor Parent Percentage | Relative Increase | Absolute Increase |
|---|---|---|---|---|
| Selous, Tanzania | 0.442 | 0.569 | 0.288 | 0.127 |
| Mikumi, Tanzania | 0.896 | 1.15 | 0.283 | 0.254 |
| Udzungwa, Tanzania | 1.23 | 1.46 | 0.189 | 0.232 |
| Ruaha, Tanzania | 7.92 | 8.15 | 0.0292 | 0.231 |
| Tarangire, Tanzania | 21.8 | 20.2 | -0.0725 | -1.58 |
| Arusha, Tanzania | 6.82 | 7.22 | 0.0585 | 0.399 |

| | | | | |
|---|---|---|---|---|
| Ngorongoro, Tanzania | 4.92 | 5.16 | 0.0491 | 0.242 |
| Gombe, Tanzania | 5.21 | 5.16 | -0.00878 | -0.0457 |
| Lake Manyara, Tanzania | 5.21 | 4.88 | -0.0633 | -0.33 |
| Serengeti, Tanzania | 3.38 | 3.34 | -0.0117 | -0.0395 |

**Supplementary Table 5:** Weighted linear regression of MPA proportion against background diversity.

| Origin | Intercept | Slope | Intercept P-value | Slope P-value | Intercept stderr | Slope stderr |
|---|---|---|---|---|---|---|
| Selous, Tanzania | 0.00263 | 0.729 | 0.000225 | 0.0404 | 0.000714 | 0.356 |
| Mikumi, Tanzania | 0.00646 | 2.19 | 6.05e-34 | 1.21e-16 | 0.000532 | 0.265 |
| Udzungwa, Tanzania | 0.00692 | 3.79 | 5.47e-15 | 9.51e-18 | 0.000886 | 0.441 |
| Ruaha, Tanzania | 0.0641 | 9.25 | 2.24e-212 | 1.91e-19 | 0.00206 | 1.03 |
| Tarangire, Tanzania | 0.211 | -0.784 | 0 | 0.663 | 0.00361 | 1.8 |
| Arusha, Tanzania | 0.0686 | 2.34 | 1.54e-166 | 0.0591 | 0.00249 | 1.24 |
| Ngorongoro, Tanzania | 0.0385 | 7.33 | 1.9e-80 | 4.03e-13 | 0.00203 | 1.01 |
| Gombe, Tanzania | 0.0306 | 10.4 | 7.72e-71 | 4.35e-34 | 0.00172 | 0.858 |
| Lake Manyara, Tanzania | 0.0408 | 4.84 | 4.67e-141 | 1.76e-09 | 0.00161 | 0.804 |
| Serengeti, Tanzania | 0.0255 | 4.4 | 2.97e-86 | 9.41e-12 | 0.0013 | 0.646 |

**Supplementary Table 6:** Minor Parent Percentage in the lowest and highest 20% Background Diversity quantile, as well as the relative and absolute difference on the autosomes.

| Origin | 0-20 Minor Parent Percentage | 80-100 Minor Parent Percentage | Relative Increase | Absolute Increase |
|---|---|---|---|---|
| Selous, Tanzania | 0.474 | 0.489 | 0.0323 | 0.0153 |
| Mikumi, Tanzania | 0.988 | 1.26 | 0.279 | 0.276 |
| Udzungwa, Tanzania | 1.22 | 1.76 | 0.443 | 0.541 |
| Ruaha, Tanzania | 7.68 | 8.92 | 0.162 | 1.25 |
| Tarangire, Tanzania | 22.4 | 21 | -0.0609 | -1.36 |
| Arusha, Tanzania | 7.13 | 7.39 | 0.036 | 0.257 |
| Ngorongoro, Tanzania | 5.14 | 5.65 | 0.1 | 0.515 |
| Gombe, Tanzania | 4.46 | 5.74 | 0.288 | 1.28 |
| Lake Manyara, Tanzania | 4.79 | 5.26 | 0.0966 | 0.463 |
| Serengeti, Tanzania | 3.23 | 3.56 | 0.103 | 0.333 |

**Supplementary Table 7:** Slope and p-value results for a GLM of the form Minor Parent Percentage ~ Recombination + Background Diversity for the autosome.

| Origin | Recombination Slope | Diversity Slope | Recombination p-val | Diversity p-val |
|---|---|---|---|---|
| Selous, Tanzania | 0.0182 | -0.794 | 0.000344 | 0.0838 |
| Mikumi, Tanzania | 0.00743 | 1.12 | 0.0592 | 0.00181 |
| Udzungwa, Tanzania | -0.00319 | 3.56 | 0.612 | 3.4e-10 |
| Ruaha, Tanzania | -0.0559 | 10.5 | 0.000113 | 1.4e-15 |

| | | | | |
|---|---|---|---|---|
| Tarangire, Tanzania | -0.0618 | -3.15 | 0.0157 | 0.173 |
| Arusha, Tanzania | -0.0215 | 2.56 | 0.223 | 0.11 |
| Ngorongoro, Tanzania | -0.0277 | 6.1 | 0.0528 | 2.48e-06 |
| Gombe, Tanzania | -0.0619 | 11.5 | 2.43e-07 | 4.45e-26 |
| Lake Manyara, Tanzania | -0.0582 | 6.27 | 2.78e-07 | 9.22e-10 |
| Serengeti, Tanzania | -0.0449 | 5.69 | 6.86e-07 | 3.43e-12 |

**Supplementary Table 8:** Diversity statistics for all 15 populations studied. Pool-Nielsen ratio is described in [4] and is a demographic adjustment for the fact that chromosome X loses diversity quicker when effective population size contracts.

| Origin | Autosomal Diversity | Pool-Nielsen Ratio | Expected chrX Diversity | Actual ChrX Diversity | P-value | Depletion Percentage | Baboon Species |
|---|---|---|---|---|---|---|---|
| Selous, Tanzania | 0.00261 | 0.739 | 0.00156 | 0.00123 | 9.22e-72 | 19.2 | yellow |
| Mikumi, Tanzania | 0.00257 | 0.745 | 0.00154 | 0.00111 | 2.14e-146 | 26.3 | yellow |
| Udzungwa, Tanzania | 0.00263 | 0.748 | 0.00159 | 0.00137 | 2.95e-41 | 11.8 | yellow |
| Ruaha, Tanzania | 0.00283 | 0.757 | 0.00173 | 0.00141 | 5.1e-74 | 16.7 | yellow |
| Tarangire, Tanzania | 0.00214 | 0.7 | 0.00121 | 0.00104 | 3.98e-34 | 12.2 | olive |
| Arusha, Tanzania | 0.0018 | 0.677 | 0.000985 | 0.000749 | 4.51e-82 | 22.3 | olive |
| Ngorongoro, Tanzania | 0.00175 | 0.653 | 0.000921 | 0.000634 | 1.23e-146 | 29.7 | olive |

| | | | | | | | |
|---|---|---|---|---|---|---|---|
| Gombe, Tanzania | 0.00164 | 0.676 | 0.000894 | 0.00058 | 1.96e-176 | 33.7 | olive |
| Lake Manyara, Tanzania | 0.00179 | 0.677 | 0.000979 | 0.000684 | 9.26e-156 | 28.6 | olive |
| Serengeti, Tanzania | 0.00182 | 0.69 | 0.00101 | 0.000667 | 3.95e-200 | 32.7 | olive |
| Gog, Ethiopia | 0.0017 | 0.679 | 0.000933 | 0.000604 | 2.29e-202 | 33.8 | olive |
| Niokolo-Koba, Senegal | 0.000509 | 0.507 | 0.000208 | 0.000176 | 1.52e-21 | 13.7 | guinea |
| Filoha, Ethiopia | 0.00166 | 0.664 | 0.000889 | 0.000713 | 8.95e-61 | 18 | hamadryas |
| Dendro Park, Zambia | 0.002 | 0.698 | 0.00113 | 0.000715 | 1.42e-176 | 35.2 | chacma |
| Chunga, Zambia | 0.00273 | 0.753 | 0.00166 | 0.00103 | 8.73e-293 | 36.6 | kinda |

**Supplementary Table 9:** Results for the weighted linear regression on chromosome X for background diversity and MPA.

| Origin | Intercept | Slope | Intercept P-value | Slope P-value | Intercept stderr | Slope stderr |
|---|---|---|---|---|---|---|
| Selous, Tanzania | 0.0605 | -33.1 | 2.57e-08 | 0.0106 | 0.0109 | 13 |
| Mikumi, Tanzania | 0.0505 | 1.64 | 1.14e-06 | 0.895 | 0.0104 | 12.4 |

| | | | | | | |
|---|---|---|---|---|---|---|
| Udzungwa, Tanzania | 0.0489 | 11.9 | 8.84e-06 | 0.366 | 0.011 | 13.1 |
| Ruaha, Tanzania | 0.031 | 156 | 0.0502 | 1.79e-16 | 0.0158 | 18.9 |
| Tarangire, Tanzania | 0.28 | -40.3 | 4.35e-53 | 0.0643 | 0.0183 | 21.8 |
| Arusha, Tanzania | 0.0399 | 32.1 | 0.00236 | 0.0403 | 0.0131 | 15.7 |
| Ngorongoro, Tanzania | -0.012 | 81.1 | 0.308 | 8.19e-09 | 0.0118 | 14.1 |
| Gombe, Tanzania | 0.0188 | 61.7 | 0.108 | 9.83e-06 | 0.0117 | 14 |
| Lake Manyara, Tanzania | 0.00314 | 65.6 | 0.772 | 3.69e-07 | 0.0108 | 12.9 |
| Serengeti, Tanzania | -0.0112 | 68 | 0.273 | 2.47e-08 | 0.0102 | 12.2 |

**Supplementary Table 10:** Results for the weighted linear regression on chromosome X for recombination rate and MPA.

| Origin | Interce pt | Slope | Intercept P-value | Slope P-value | Intercept stderr | Slope stderr |
|---|---|---|---|---|---|---|
| Selous, Tanzania | 0.0413 | -0.075 | 1.8e-09 | 0.258 | 0.00686 | 0.0664 |
| Mikumi, Tanzania | 0.0457 | 0.075 1 | 2.91e-12 | 0.235 | 0.00654 | 0.0633 |
| Udzungwa, Tanzania | 0.0479 | 0.122 | 4.06e-12 | 0.0672 | 0.0069 | 0.0668 |

| | | | | | | |
|---|---|---|---|---|---|---|
| Ruaha, Tanzania | 0.0961 | 0.658 | 5.88e-22 | 9.23e-12 | 0.00997 | 0.0965 |
| Tarangire, Tanzania | 0.255 | -0.0716 | 1.99e-109 | 0.519 | 0.0115 | 0.111 |
| Arusha, Tanzania | 0.0479 | 0.201 | 6.34e-09 | 0.0118 | 0.00826 | 0.0798 |
| Ngorongoro, Tanzania | 0.0235 | 0.337 | 0.00164 | 3.04e-06 | 0.00746 | 0.0722 |
| Gombe, Tanzania | 0.0495 | 0.197 | 1.93e-11 | 0.00562 | 0.00737 | 0.0713 |
| Lake Manyara, Tanzania | 0.0439 | 0.126 | 1.7e-10 | 0.0574 | 0.00687 | 0.0664 |
| Serengeti, Tanzania | 0.0174 | 0.29 | 0.00695 | 3.57e-06 | 0.00646 | 0.0625 |

**Supplementary Table 11:** Minor Parent Percentage in the lowest and highest 20% Background Diversity quantile for chromosome X, as well as the relative and absolute difference on the autosomes.

| Origin | 0-20 Minor Parent Percentage | 80-100 Minor Parent Percentage | Relative Increase | Absolute Increase |
|---|---|---|---|---|
| Selous, Tanzania | 6.56 | 2.54 | -0.613 | -4.02 |
| Mikumi, Tanzania | 6.45 | 5.14 | -0.203 | -1.31 |
| Udzungwa, Tanzania | 6.76 | 5.97 | -0.118 | -0.797 |
| Ruaha, Tanzania | 7.94 | 20.1 | 1.54 | 12.2 |
| Tarangire, Tanzania | 25.7 | 23.6 | -0.0806 | -2.07 |
| Arusha, Tanzania | 1.21 | 7.25 | 4.98 | 6.04 |

| | | | | |
|---|---|---|---|---|
| Ngorongoro, Tanzania | 0.471 | 7.27 | 14.4 | 6.8 |
| Gombe, Tanzania | 4.3 | 8.49 | 0.975 | 4.19 |
| Lake Manyara, Tanzania | 1.31 | 7.1 | 4.44 | 5.8 |
| Serengeti, Tanzania | 0.704 | 6.37 | 8.04 | 5.66 |

**Supplementary Table 12)** Interaction Slope denotes the difference between the slopes of the normalized regression based on autosomes and chromosome X, using background diversity as the explanatory variable.

| Origin | Autosome Slope | Interaction Slope | Relative Increase | Autosome P-value | Interaction Slope P-value |
|---|---|---|---|---|---|
| Selous, Tanzania | 0.000429 | -0.0102 | -23.7 | 0.0626 | 2.72e-08 |
| Mikumi, Tanzania | 0.00123 | -0.000699 | -0.568 | 1.35e-11 | 0.626 |
| Udzungwa, Tanzania | 0.00225 | 0.00165 | 0.733 | 1.65e-16 | 0.458 |
| Ruaha, Tanzania | 0.00536 | 0.0382 | 7.12 | 1.41e-18 | 1.42e-14 |
| Tarangire, Tanzania | -0.00101 | -0.00849 | 8.41 | 0.337 | 0.322 |
| Arusha, Tanzania | 0.0013 | 0.00952 | 7.32 | 0.0742 | 0.109 |

| | | | | | |
|---|---|---|---|---|---|
| Ngorongoro, Tanzania | 0.00395 | 0.0218 | 5.53 | 2.86e-11 | 6.29e-06 |
| Gombe, Tanzania | 0.00604 | 0.0125 | 2.08 | 7.84e-33 | 0.00235 |
| Lake Manyara, Tanzania | 0.00272 | 0.0182 | 6.68 | 9.95e-09 | 2.53e-06 |
| Serengeti, Tanzania | 0.00265 | 0.0189 | 7.13 | 4.83e-12 | 1.42e-09 |

**Supplementary Table 13:** MPA proportion across autosomes and chromosome X for all investigated populations. Every chromosomes MPA proportion is taken as one observation, weighting them equally.

| Origin | Mean Autosome MPA | Mean ChrX MPA | Mann-Whitney P-value | Autosome - chrX |
|---|---|---|---|---|
| Selous, Tanzania | 0.414 | 4.52 | 8.06e-05 | -4.11 |
| Mikumi, Tanzania | 1.09 | 5.72 | 1.06e-64 | -4.63 |
| Udzungwa, Tanzania | 1.38 | 6.14 | 2.17e-08 | -4.75 |
| Ruaha, Tanzania | 8.17 | 12.9 | 1.96e-05 | -4.74 |
| Tarangire, Tanzania | 21 | 25 | 0.364 | -3.96 |
| Arusha, Tanzania | 7.19 | 6.2 | 0.678 | 0.99 |
| Ngorongoro, Tanzania | 5.28 | 4.2 | 0.0268 | 1.08 |
| Gombe, Tanzania | 4.99 | 6.31 | 0.0104 | -1.31 |
| Lake Manyara, Tanzania | 4.93 | 5.05 | 0.534 | -0.123 |
| Serengeti, Tanzania | 3.41 | 3.48 | 0.756 | -0.0705 |

**Supplementary Table 14:** Autosome/Chromosome X regression comparison after filtering out windows with more than 25 % MPA. Interaction Slope denotes the difference between the slopes of the normalized autosomal regression and chrX.

| Origin | Autosome Slope | Interaction Slope | Relative Increase | Autosome P-value | Interaction Slope P-value |
|---|---|---|---|---|---|
| Selous, Tanzania | 8.82e-05 | 0.00176 | 20 | 0.491 | 0.091 |
| Mikumi, Tanzania | 0.00137 | 0.00893 | 6.51 | 6.63e-22 | 7.26e-15 |
| Udzungwa, Tanzania | 0.00216 | 0.00791 | 3.66 | 1.63e-22 | 2.35e-05 |
| Ruaha, Tanzania | 0.00277 | 0.0174 | 6.3 | 7.91e-10 | 1.98e-05 |
| Tarangire, Tanzania | -0.00144 | 0.0025 | -1.74 | 0.0175 | 0.622 |
| Arusha, Tanzania | 0.000176 | 0.00121 | 6.92 | 0.711 | 0.75 |
| Ngorongoro, Tanzania | 0.00173 | 0.00252 | 1.46 | 5.99e-06 | 0.426 |
| Gombe, Tanzania | 0.00331 | 0.00243 | 0.733 | 4.85e-21 | 0.402 |
| Lake Manyara, Tanzania | 0.00131 | 0.0101 | 7.67 | 0.000117 | 0.000309 |
| Serengeti, Tanzania | 0.0015 | 0.00715 | 4.77 | 2.82e-07 | 0.00316 |

**Supplementary Table 15:** Slope and p-value results for a GLM of the four combinations of Autosome/chrX and Recombination/Expected Diversity for Gog Olives when inferring admixture from Hamadryas.

| Autosomes | Intercept | Slope | Intercept P-value | Slope P-value | Intercept stderr | Slope stderr |
|---|---|---|---|---|---|---|
| Gog, Ethiopia, Recombination | 0.0658 | 0.251 | 9.72e-272 | 1.03e-76 | 0.00187 | 0.0136 |
| Gog, Ethiopia, Diversity | 0.0197 | 38.8 | 5.74e-10 | 1.58e-132 | 0.00318 | 1.58 |
| **ChrX** | | | | | | |
| Gog, Ethiopia, Recombination | 0.279 | 1.07 | 1.45e-52 | 1.39e-09 | 0.0183 | 0.177 |
| Gog, Ethiopia, Diversity | 0.205 | 214 | 1.85e-12 | 7.18e-10 | 0.0291 | 34.7 |

#### References


1. Maples BK, Gravel S, Kenny EE, Bustamante CD. RFMix: A Discriminative Modeling Approach for Rapid and Robust Local-Ancestry Inference. Am J Hum Genetics. 2013;93:278–88.
2. Massarat AR, Lamkin M, Reeve C, Williams AL, D'Antonio M, Gymrek M. Haptools: a toolkit for admixture and haplotype analysis. Bioinformatics. 2023;39:btad104.
3. Kamm JA, Spence JP, Chan J, Song YS. Two-Locus Likelihoods Under Variable Population Size and Fine-Scale Recombination Rate Estimation. Genetics. 2016;203:1381–99.
4. Pool JE, Nielsen R. POPULATION SIZE CHANGES RESHAPE GENOMIC PATTERNS OF DIVERSITY. Evolution. 2007;61:3001–6.
